# Spatio-temporal patterning of living cells with extracellular DNA programs

**DOI:** 10.1101/2020.05.14.096065

**Authors:** Marc Van Der Hofstadt, Jean-Christophe Galas, André Estevez-Torres

## Abstract

Reactive extracellular media focus on engineering reaction networks outside the cell to control intracellular chemical composition across time and space. However, current implementations lack the feedback loops and out-of-equilibrium molecular dynamics for encoding spatio-temporal control. Here, we demonstrate that enzyme-DNA molecular programs combining these qualities are functional in an extracellular medium where human cells can grow. With this approach, we construct an internalization program that delivers fluorescent DNA inside living cells and remains functional for at least 48 h. Its non-equilibrium dynamics allows us to control both the time and position of cell internalization. In particular, a spatially inhomogeneous version of this program generates a tunable reaction-diffusion two-band pattern of cell internalization. This demonstrates that a synthetic extracellular program can provide temporal and positional information to living cells, emulating archetypal mechanisms observed during embryo development. We foresee that non-equilibrium reactive extracellular media could be advantageously applied to *in vitro* biomolecular tracking, tissue engineering or smart bandages.

---

Traditionally, synthetic biology has focused its engineering efforts inside the cells, through the assembly of relatively complex *intracellular* genetic circuits.^1^ These circuits have been driven from the outside by using simple extracellular inputs (such as small molecules^2,3^ or light^4^), resulting, for instance, in the control of cellular fate^5^ and intracellular composition.^6^ A complementary route consists in designing a reactive *extracellular* medium to control the intracellular composition. Such strategy could be advantageously applied in tissue engineering^7^ or smart bandages,^8^ but also to better understand cells’ behaviour in a changing environment.^9^

The reactive extracellular approach has been explored in the last decade with the tools of DNA nanotechnology and molecular programming.^10^ Indeed, extracellular DNA nanostructures have been used for delivering payloads *in vitro*^11^ and *in vivo*,^12^ for controlling cell adhesion, ^13^ and for intracellular detection. ^14^ In addition, nucleic acid molecular programs are able to classify cells depending on the biomarkers displayed on their surface^15–17^ and are functional inside cells. ^18,19^ However, to our knowledge, these methods have not yet been able to dynamically control the composition of the extracellular medium in the presence of viable cells.

The design of reactive extracellular media has been pioneered by Mansy and co-workers using transcription-translation (TX-TL) reaction networks.^20–23^ In this approach, the extracellular reactive medium is protected from the cells inside liposomes and the interaction between the two happens through small molecules carried by transmembrane channels. The strength of this approach relies on the diversity of small molecules that may be used to entertain the conversation between synthetic and living cells. The limitation, in turn, is the relatively simple reaction dynamics that can be implemented by cell-free TX-TL networks. This is due to the rapid consumption of energy in the translation step, which rapidly drives the reaction to chemical equilibrium in a closed reactor. For this reason, out-of-equilibrium TX-TL reaction networks with feedback loops have only been observed in open reactors^24,25^ without cells. However, such non-equilibrium networks are key elements contributing to the complexity of living cells. They encode important behaviors such as bistability, homeostasis, temporal control and pattern formation but, to our knowledge, have not yet been observed in reactive extracellular media.

In this work, we have engineered a reactive medium that can support out-of-equilibrium reaction networks with feedbacks in the presence of living human cells. The reactive medium is based on DNA-enzyme molecular programs that autonomously control the concentration of selected chemicals which in turn modify the intracellular composition of living cells, both being regulated in time and space. Firstly, we demonstrate that an autocatalytic amplifier and a bistable switch are active and remain out of equilibrium for tens of hours in the presence of living human cells. In particular, this means that extracellular DNA can be detected among viable cells. Secondly, we show that these programs can trigger the internalization of fluorescently-labelled DNA and thus control the chemical composition of living cells. Importantly, its out-of-equilibrium dynamics allows the program to remain at steady state for 48 h and to encode a temporal control. Such temporal control can be triggered externally, by responding to external inputs, or internally, through a clock reaction that causes internalization at times that are chemically encoded. Finally, we demonstrate that a reactive extracellular medium processes the spatial information contained in a concentration gradient and transfers it to living cells, recapitulating a major mechanism of pattern formation in embryo development:^26^ a shallow extracellular gradient of DNA creates a sharp two-band pattern of fluorescent DNA that ultimately patterns living cells.

## A common buffer for DNA programs and living cells

The PEN DNA toolbox is a DNA programming framework that implements the three basic operations of regulatory networks — activation, repression and degradation — using short DNA oligonucleotides and 3 enzymes, namely a polymerase, an exonuclease and a nicking enzyme (PEN). The use of enzymes allows the implementation of non-equilibrium reaction networks with feedbacks^28,29^ that display robust^27^ dynamics, such as oscillations,^28,30^ bistability ^31,32^ and spatial patterns.^33,34^

Figure 1a depicts the core element of PEN networks used throughout this work: an autocatalytic module that exponentially amplifies an oligonucleotide **A**, before reaching a steady state. Briefly, **A** is a single-stranded DNA (ssDNA), called trigger, and **T** is a ssDNA, called template, twice complementary to **A**. Upon hybridization of **A** on the 3’ end of **T**, a polymerase enzyme elongates **A** until it forms a fully double-stranded DNA (dsDNA). A nicking enzyme cuts the elongated strand, generating two copies of **A**, that dissociate from **T**, and hence show exponential amplification through the net reaction **A** → 2**A**. Simultaneously, an exonuclease enzyme selectively degrades free **A** and, after some time, the equilibration of autocatalysis and degradation yields a steady state where the concentration of **A**, noted [A], is constant and equal to [A]_*ss*_. This non-equilibrium steady state is maintained until the deoxynucleotides (dNTPs) present in the solution are consumed, which takes several days.^35^ Our first objective was to find common experimental conditions where PEN autocatalysis was functional and cells were viable.

**Figure 1:**
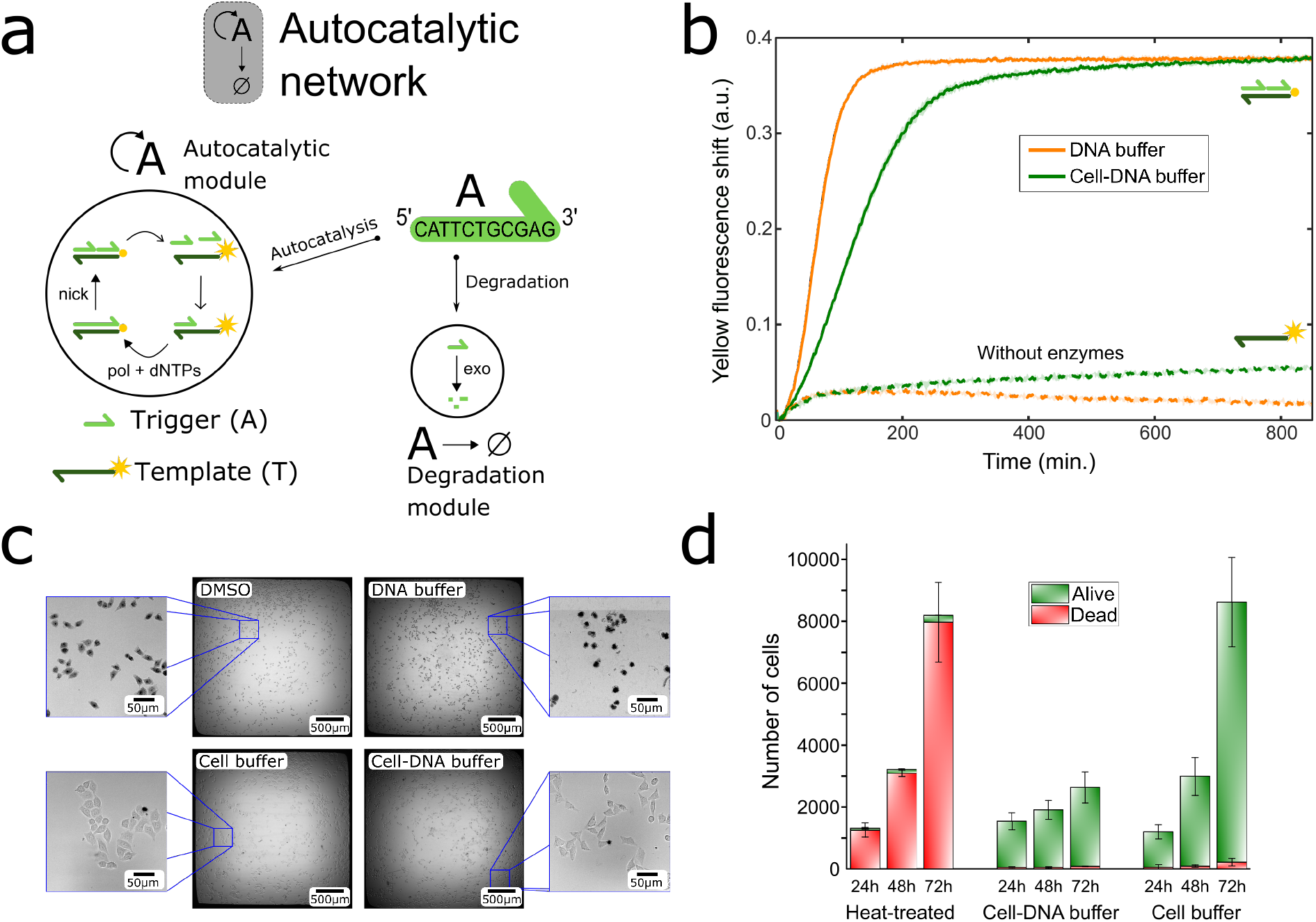
DNA autocatalysis is functional and cells are viable in a common buffer. a) Scheme of the core autocatalytic network used throughout this work, composed of an autocatalytic and a degradation module. ssDNAs are represented as harpoon-ended arrows. During autocatalysis, the trigger **A** replicates on template **T** through a combination of hybridization and dehybridization steps, elongation by a polymerase (pol), and cutting by a nicking enzyme (nick). Upon elongation of **A**, the fluorescent marker attached to **T** (yellow star) gets quenched. Specific degradation of **A** is carried out by an exonuclease enzyme (exo). Solid and empty arrow-heads indicate irreversible and reversible reactions, respectively. b) Fluorescence shift from the fluorescently-labelled **T** versus time for the autocatalytic network in standard DNA buffer (orange line), or in the biocompatible cell-DNA buffer (green line), at 37 °C in the absence of cells. Dashed lines correspond to negative controls in the absence of enzymes. The shades, corresponding to the standard deviation of a triplicate experiment, are of the order of the line thickness. c) Bright-field images of trypan blue-stained cells after 24 h incubation in different buffers, dead cells appear in black. Treatment with DMSO is a control for dead cells. The black halo around the central images is an optical artifact. d) Number of dead and live cells per well determined by propidium iodide staining and flow cytometry for different incubation times and conditions (experiment performed per triplicate and repeated 2 different days, *n* = 6). In panels c and d the cell-DNA buffer contained the autocatalytic network. Conditions: [A]_0_ = 0.5 nM, [T]_0_ = 100 nM in panel b and [A]_0_ = 20 nM, [T]_0_ = 200 nM in c and d.

The majority of PEN autocatalytic modules operate between 42-45 °C in an optimal buffer,^36^ hereafter referred as *DNA buffer*. Nevertheless, standard human cell culture conditions require 37 °C and a rich medium, such as Dulbecco’s modified Eagle’s medium (DMEM). We thus screened different buffer compositions and found a *cell-DNA buffer* compatible with both systems (SI Section 2 and Figures S1-S4). In addition, the sequences of **A** and **T** were redesigned to increase the autocatalytic efficiency at 37 °C (SI Table S2 and Figure S5).

To assess the performance of the autocatalytic network in the new buffer, we first monitored its dynamics *in the absence of cells*. To do so, strand **T** had a yellow fluorophore attached to its 5’ end that was quenched upon hybridization; the shift in fluorescence signal being proportional to [A] if [A] < [T]_0_, where [T]_0_ is the initial concentration of **T**.^31^ Figure 1b shows that autocatalytic dynamics are qualitatively similar in the DNA and the cell-DNA buffers in the absence of cells, with a sigmoidal curve characteristic of PEN autocatalytic reactions.^28^

The amplification kinetics were 2-fold slower in the cell-DNA buffer, which we attribute to a lower efficiency of the enzymes (Figure S6). As expected, no reaction was detected in the absence of PEN enzymes. Importantly, the concentration of **A** at steady state was identical in both buffers: starting from [A]_0_ = 0.5 nM, **A** reached 400 nM, a 800-fold amplification, within 200 min (Figure S5).

We then measured the viability of living human cervix epithelial carcinoma cells (HeLa cell line) in the different buffers and in the presence or in the absence of the autocatalytic network. Trypan blue cell staining after 24 h revealed large mortality of the cells in the DNA buffer, while the mortality remained low and comparable in the cell-DNA and in the reference cell-buffer (Figure 1c). However, propidium iodide cell staining combined with flow cytometry (Figure 1d) showed that cells divided ~4-fold slower in the cell-DNA buffer than in the cell-buffer (doubling time of 61 and 17 h, respectively). We account this to the presence of DTT that causes partial cell detachment (Figure S2) and transiently activates endoplasmic reticulum stress.^37^ In summary, cell growth was slower in the cell-DNA buffer but cell death was negligible for up to 3 days.

## An extracellular medium with out-of-equilibrium dynamics

A key point of PEN reactions is the possibility to engineer out-of-equilibrium reaction networks with feedback loops. To demonstrate that this remains true in a solution containing living cells, we quantified the dynamics of two such networks in the cell-DNA buffer in the presence and in the absence of cells, using the DNA-buffer without cells as a positive control (see Methods).

Autocatalysis is the most basic feedback loop and it is essential to build out-of-equilibrium dynamics.^38^ Figure 2a displays the amplification onset time, *τ*, of the autocatalytic network as a function of the initial concentration of **A**, [A]_0_. In all three cases, *τ* depends linearly on log([A]_0_), indicating that autocatalysis followed first order kinetics. In addition, a moderate reduction in the rate of the autocatalytic reaction was observed: by 20% between the DNA and the cell-DNA buffer in the absence of cells and by a further 20% in the presence of living cells. Importantly, these data show that this DNA program was able to detect a specific DNA sequence at a concentration of 100 pM in the presence of cells, within 300 min.

**Figure 2:**
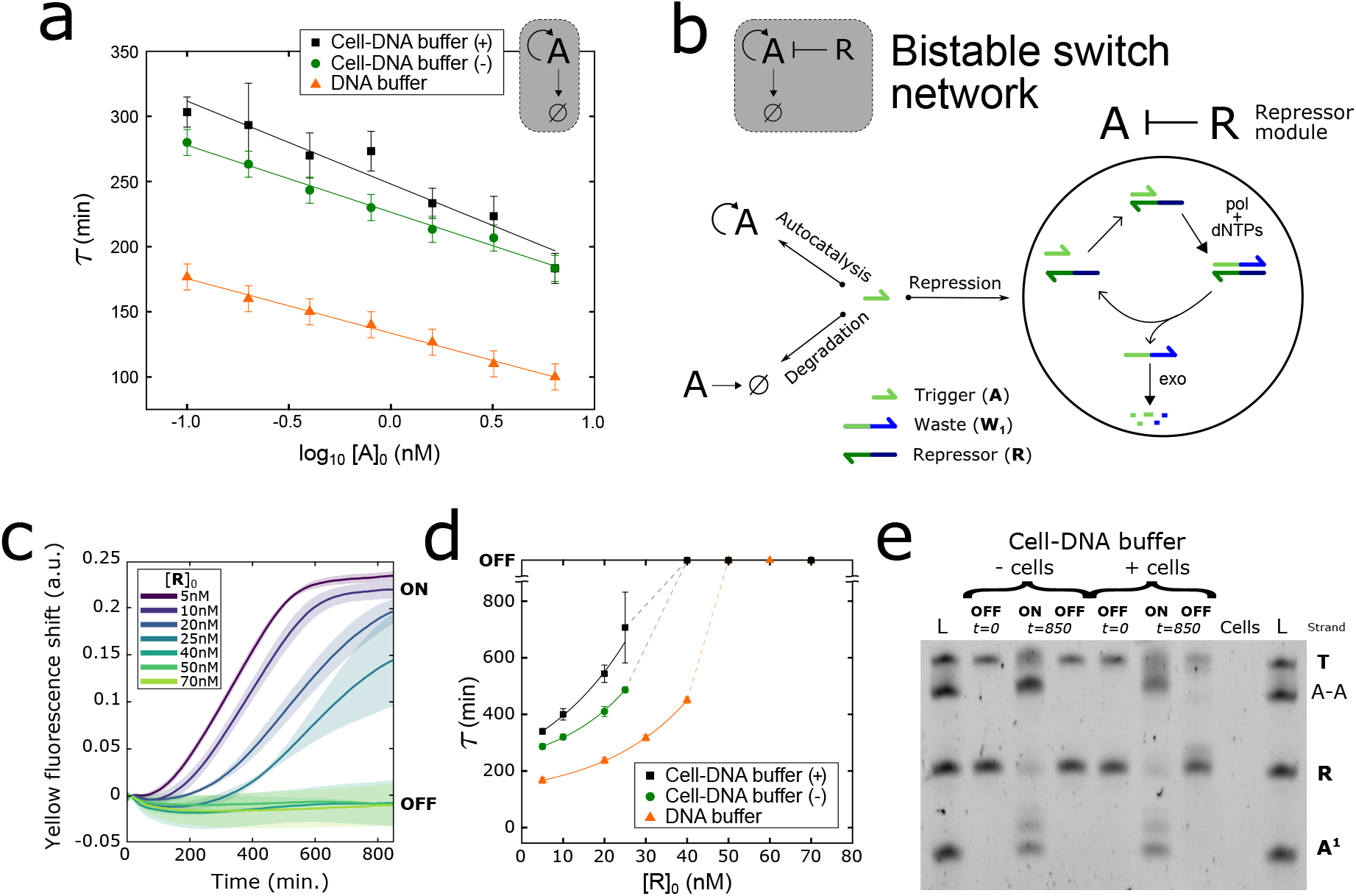
Out-of-equilibrium networks with feedbacks are fully functional in the presence of cells. a) Amplification onset time, *τ*, against the decimal logarithm of the initial concentration of **A** for the autocatalytic network in two different buffers, and in the presence ((+), black squares) or in the absence ((-), green circles) of cells. b) A bistable switch (in gray) is obtained when the autocatalytic network is combined with a repressor module, where **A** binds to repressor **R** and is converted into a non-functional ssDNA, waste **W_1_**, that is degraded by the exonuclease. c) Fluorescent shift from fluorescently-labelled **T** versus time for the bistable switch in the cell-DNA buffer in the presence of cells for increasing concentrations of **R**, [R]_0_. d) *τ* versus [R]_0_ for the bistable switch in the same conditions as in panel a. Data determined from panel c and Figure S7. e) Denaturing polyacrylamide gel electrophoresis showing the products obtained in the *OFF* and *ON* states of the bistable switch in the absence (-) or in the presence (+) of cells, at different times. L is a ladder containing the template **T** (20 nt with a JOE fluorescent label), the strand A-A (a 22 nt resulting from the elongation of **A** on **T**), repressor **R** (15 nt) and species **A**^1^ (which corresponds to species **A** with a phosphate group). The ‘cells’ lane is an extract from a solution containing cells only. Experiments were performed in 384-well cell culture plates. The error bars and the shades in panels a, c and d correspond to one standard deviation of a triplicate experiment. Solid lines in panels a and d are linear and exponential fits, respectively, while dashed lines are guides to the eye. Conditions: [T]_0_ = 100 nM in all panels, and [A]_0_ = 0.5 nM in panels c and d.

Chemical bistability is intrinsically a non-equilibrium property. ^38^ The PEN autocatalytic network can be reprogrammed into a bistable switch^32^ by adding a repressor strand **R**, creating a reactive system with two antagonistic steady-states: an *ON* state where autocatalysis of **A** occurs and [A]_*ss*_ is high, and an *OFF* state, where the combined degradation of **A** by **R** and the exonuclease outcompetes autocatalysis, yielding [A]_*ss*_ = 0 (Figure 2b). Figure 2c shows the dynamics of the bistable switch in the presence of cells as a function of the concentration of **R**, at constant [A]_0_. Increasing [R]_0_ slows down the exponential amplification of **A** until the system switches to the *OFF* state at [R]_0_ > 40 nM, where no amplification was detectable within 850 min. The plot of *τ* as a function of [R]_0_ highlights a strong non-linear response that is characteristic of a bifurcation point (Figure 2d). Again, the presence of the cells does not qualitatively change the dynamics of the bistable switch in the cell-DNA buffer. Gel electrophoresis analysis confirmed that the bistable switch produced the same products in the presence and in the absence of cells (Figure 2e), with a similar pattern to the one previously obtained in standard conditions.^27^

## A reactive medium that controls cellular internalization

Until now we have shown the coexistence of two orthogonal out-of-equilibrium systems: the cells and the DNA programs. We now explore the possibility of engineering out-of-equilibrium DNA programs that alter the chemical composition of living cells. To do so, we created an internalization switch by coupling the bistable switch to the conversion module **A** → **S**, that releases fluorescently-labeled ssDNA signal **S**, later internalized by the cells (Figure 3a). The conversion module relies on the reaction **A** + **C** → **S** + **W**_2_, where complex **C** is a partially double-stranded DNA whose shorter top strand is **S** and whose longer bottom strand (**D**) is attached to a hydrogel bead to avoid uncontrolled internalization, hereafter termed conversion bead. **A** is designed to bind to the bottom strand of complex **C** and release the top strand **S** by polymerase-assisted strand displacement into the extracellular medium. Once in solution, **S** is passively internalized by living cells^39^ within 2–4 hours, making them fluorescent (SI Video S1 and Figure S8). Figure 3b sketches the experimental setup, where the cells and the conversion beads were placed in the bottom of a cell culture well filled with a reactive medium containing the cell-DNA buffer and the bistable switch. The beads were homogeneously distributed among the well’s bottom and covered 2.6% of its surface (SI Figure S9).

**Figure 3:**
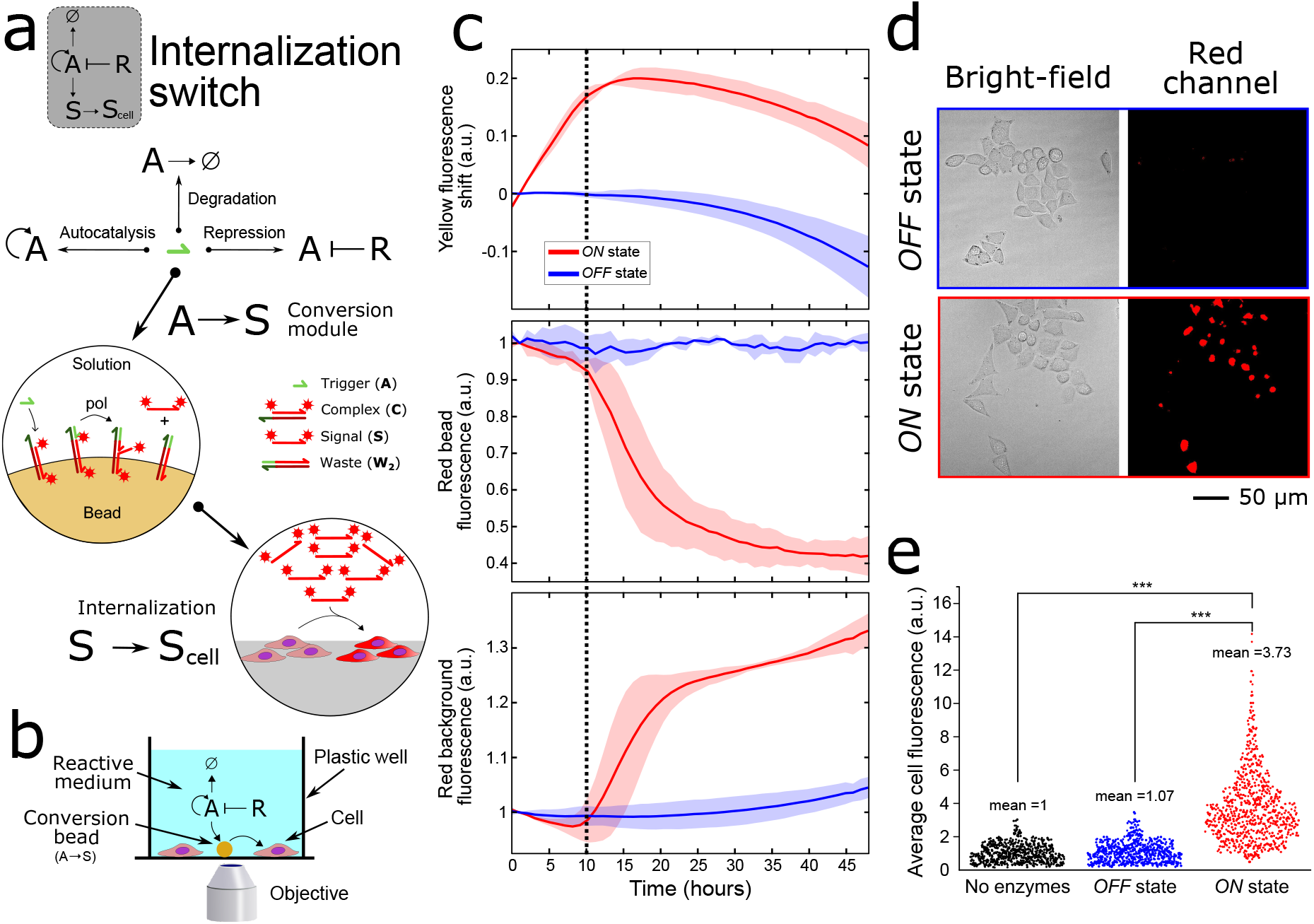
A reactive medium that controls the internalization of fluorescent DNA inside living cells. a) Scheme describing the operation of the DNA internalization switch embedded in the reactive medium. The gray box depicts the topology of the reaction network, composed of a bistable switch and a conversion module. In the conversion module, free **A** binds to complex **C**, attached to hydrogel beads, and is elongated by the polymerase. This causes the release of the red fluorescent strand **S** into the solution, that can be passively internalized by the cells (**S**_*cell*_). Red stars indicate attached red fluorophores. b) Cartoon of the experimental setup (not to scale). The cells are cultured in a reactive medium containing the bistable switch in solution and the conversion module attached to hydrogel beads of 34 *μ*m diameter. c) Fluorescence versus time plots displaying the dynamics of the internalization switch in the *ON* (red line) and *OFF* (blue line) states in the presence of cells: production of **A** (top) and release of **S** from the beads (middle) into the solution (bottom). The black dashed line is a guide to the eye. d) Overlapping bright-field and red fluorescence microscopy images of living cells rinsed after being cultured for 48 h in the reactive medium in the *OFF* and *ON* state. e) Quantitative analysis of the average red fluorescence intensity per cell for cells cultured in the reactive medium in the absence of enzymes (black dots), in the *OFF* state (blue dots), and in the *ON* state (red dots). At least 649 cells were analysed in each condition. Experiments were performed in 384-well culture plates in duplicates and repeated 3 different days (6 experiments in total). The shade in panel c corresponds to one standard deviation. Conditions: [A]_0_ = 20 nM, [T]_0_ = 200 nM. The *ON* and *OFF* states correspond to [R]_0_ = 0 and 200 nM, respectively.

The conversion beads are reminiscent of the liposomes used by Mansy and coworkers to protect TX-TL networks. ^20,22^ However, here only the signal strand is physically separated from the cells while the other reactive components freely float in the medium. PEN reactions on hydrogel beads where first described by Gines et al. ^40^ Note that our implementation is slightly different as it concerns a polymerase-assisted strand-displacement reaction which is a combination of PEN with DNA strand displacement^41^ reactions.

The conversion module was designed such that **A** hybridized to **C** less favourably than both to **T** and **R**, and thus **S** was released only when the bistable switch was *ON* and [A] was high (SI Figure S10). This second non-linearity makes the network particularly robust, as we will see below. Figures 3c and S11 show the dynamics of production of **A** and release of **S** in the medium for the internalization switch in the presence of cells in the *ON* and *OFF* states. In the *ON* state, **A** is amplified during at least 10 h, when it saturates **T** and causes its fluorescent signal to reach a maximum before decreasing slowly. This indicates that at *t* =10 h, [A]≈ 2[T]_0_ = 400 nM in the medium, a 20-fold amplification. At this point, there was enough free **A** to trigger the conversion reaction, releasing **S** from the beads into the extracellular medium. This process resulted in a decrease and an increase in the red fluorescence of the conversion beads and the medium, respectively, and took another 10 h. In contrast, in the *OFF* state, no amplification of **A** was observed and the release of **S** in the medium was negligible after 20 h and was at least 8-fold lower after 48 h. Note that the initial concentration of **A** (20 nM) would be sufficient for the polymerase to convert the total quantity of **C** in the well (18 nM) into **S**, but this release did not occur until [A] was high due to the non-linearity in the conversion module described above. Moreover, the evaporation of the medium was significant during this 48 h-long experiment (30%). This could explain the long-term decrease and increase of the fluorescent signals used to follow **A** and **S** in the medium (SI Figure S12) but did not compromise the function of the DNA program.

After 48 h, the cells were rinsed to eliminate the excess of free **S** in the medium and imaged. Figure 3d shows that only the cells cultured in the presence of the internalization switch in the *ON* state appear fluorescent. Higher magnification images (SI Figure S13) revealed an internalization into dot and rod-like compartments, which is compatible with an accumulation in mitochondria as observed for cyanine fluorophores^42^ similar to the one attached to **S**. We observed that cells became fluorescent only when **S** was labelled with certain fluorophores and not others (Figure S14). Figure 3e displays the distribution of the average fluorescent intensity per cell, demonstrating that the *ON* state results in a rate of cell internalization significantly different from the *OFF* state (two-tailed Mann-Whitney U test, p-value <0.001), with, respectively, means of 3.7 and 1.1, medians of 3.3 and 1.0 and standard deviations of 2.13 and 0.62. A negative control in the absence of enzymes produced a distribution of cellular fluorescence not significantly different to the *OFF* state, showing that the concentration of the red fluorophore inside the cells is controlled by the DNA program.

## The extracellular medium controls the timing of internalization

Since the bistable switch is maintained out of equilibrium, it should be possible to turn its *OFF* state *ON* and hence trigger internalization at different times. This timing could be either externally triggered or pre-programmed in the reactive medium.

To test triggered internalization, we added either a ssDNA activator **R**^*^, or a random sequence to the *OFF* state. **R**^*^ is complementary to **R** and, upon hybridization, it reduces [R], and hence switches the system to the *ON* state. Note that **R**^*^ binds in a reversible manner to **R** at 37 °C, so we used a 4.5-fold excess of **R**^*^ to shift the equilibrium towards the hybridization of **R**^*^ with **R**. Figures 4a-b and S15 show the response of the internalization switch when **R**^*^ or a random sequence was introduced 6 h after the beginning of the experiment, together with controls for the *ON* and *OFF* states. The switch worked as expected. No significant difference was observed in the *OFF* state after the addition of the random sequence. In contrast, the addition of **R**^*^ switched on the autocatalysis, which further caused the release of **S**, with delays of 16 h and 13 h, respectively, compared to the *ON* state. As anticipated, only the cells where the DNA activator was introduced internalized the fluorescent strand **S** (Figure 4b) (two-tailed Mann-Whitney U test, p-value <0.001), showing that the reactive extracellular medium was responsive for at least 6 h, and was able to change cellular composition for up to 48 h of cell culture.

**Figure 4:**
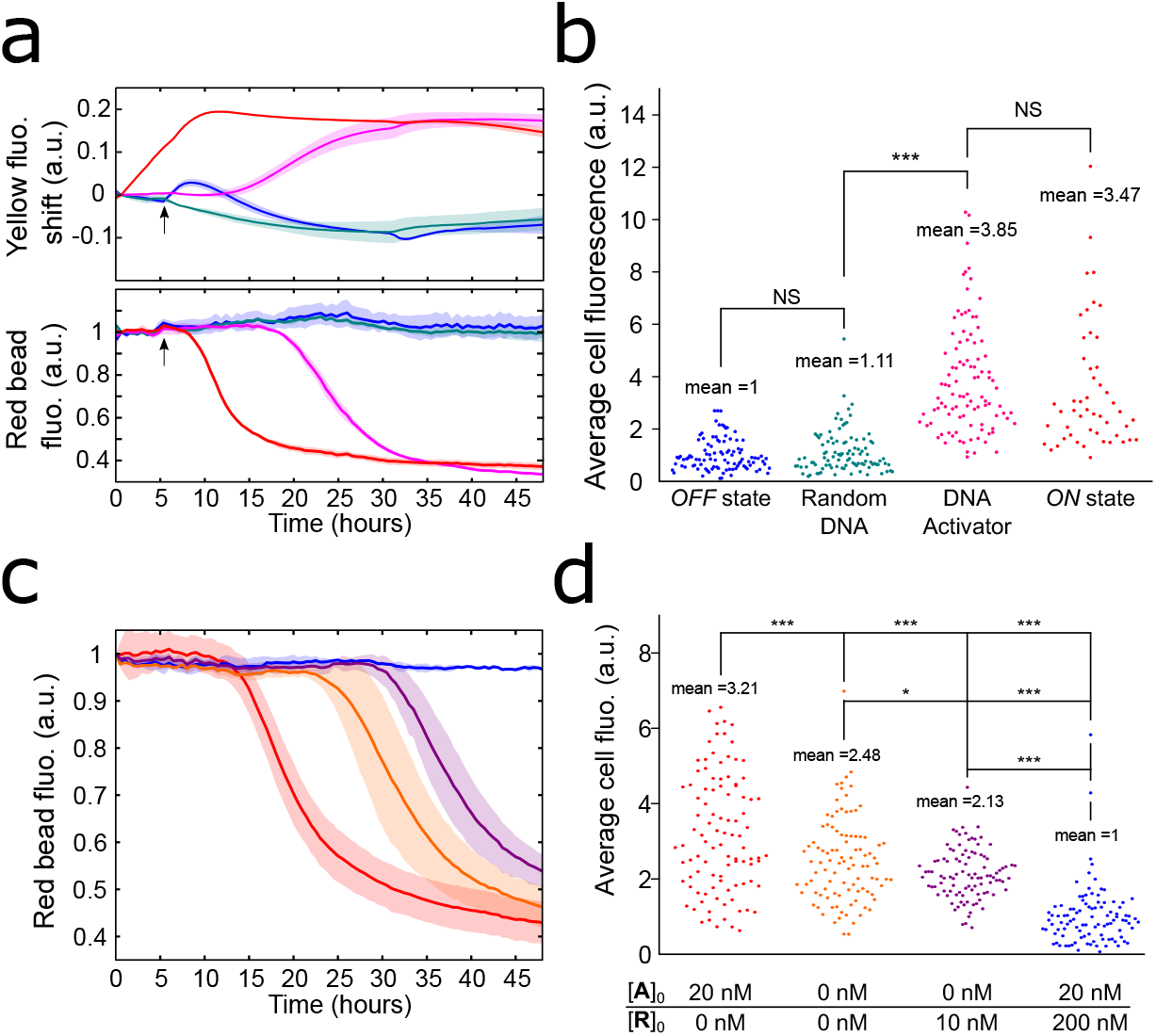
The extracellular medium controls the internalization time and the amount of **S** delivered to the cells. a) Fluorescence versus time plots showing the dynamics of production of **A** (top) and the release of **S** from the beads (bottom) in the presence of cells. The curves show the unperturbed *ON* (red) and *OFF* (blue) states, as well as *OFF* states where a ssDNA, either **R**\* (pink) or a random sequence (teal), was introduced at t = 6 h (indicated with an arrow). The bump is an artifact of the addition step. b) Quantitative analysis of the average red fluorescence intensity of living cells after 48 h exposure to the conditions in panel a (n = 50). Fluorescence versus time plots showing the release of **S** from the beads (c) and the resulting average red fluorescence intensity of living cells after 48 h (d) for different preprogrammed dynamics of the reactive medium. Each color corresponds to different initial concentrations of **A** and **S** as indicated in the bottom of panel d. 100 cells were measured for each condition in panel d. The shade in panels a and c correspond to one standard deviation of a duplicate experiment performed on the same day. NS: Not significant. Conditions for panel a and b: [A]_0_ = 20 nM, [T]_0_ = 200 nM. The *ON* and *OFF* states correspond to [R]_0_ = 0 and 200 nM, respectively and ssDNAs were added at 900 nM. Conditions for panel c and d: [T]_0_ = 200 nM.

The bistable switch can also be used as a clock reaction that triggers the release of **S** at a pre-encoded time. To do so we tuned the kinetics of destabilization of the *OFF* state by changing the initial concentrations of **A** and **R** (Figures 4c and d, and S16). Depending on the initial conditions, the timing of the release of **S** from the beads could be adjusted between 13 and 30 h. Importantly, this delay was controlled independently from the kinetics of the release step, that remained unchanged. By delaying the release, we expect to decrease the effective concentration of the red fluorescent strand **S** at which the cells are exposed. Figure 4d shows that the higher the delay of the fluorescent release, the lower the average cell fluorescence measured after 48 h (two-tailed Mann-Whitney U test, p-value <0.05 or <0.001), indicating that the amount of **S** inside the cell can be pre-encoded in the reactive medium.

## The extracellular medium provides positional information to the cells

Another characteristic property of out-of-equilibrium reactions with feedbacks is their ability to give rise to spatial concentration patterns via reaction-diffusion mechanisms. ^38^ In this regard, PEN reactions are known to display a rich variety of spatial patterns of DNA concentration.^33,34,40,43^ In particular, a bistable switch in the presence of a gradient of repressor creates a non-equilibrium two- or three-band pattern at steady state. ^34^ This last process emulates an important mechanism of pattern formation observed during early embryo development, and called positional information.^26^ During this mechanism, a shallow morphogen gradient along one axis of the embryo is translated by a gene regulatory network into a sharp pattern of protein concentration, known as the French flag. ^44^ This protein pattern is essential and widely conserved in eukaryotic organisms for the development of the anterior-posterior axis of the embryo during cell differentiation.^45^

To test whether patterning by positional information was possible in reactive extracellular media, we cultured the cells in a PDMS/glass millifluidic channel in the presence of the internalization switch (Figure 5a). The channel was filled in such a way that the composition of the medium was homogeneous along the channel, except for **R** that formed a gradient from 0 to 2 *μ*M, with high concentration on the right-hand-side. Figure 5b displays fluorescence profiles along the channel at an initial time for four DNA species that confirm this expectation, except for a slight inhomogeneity for species **A**. Indeed, the first quantitative profile was recorded 1 h after the assembly of the reactive medium and thus **A** can already be detected where the concentration of **R** is low because autocatalysis starts right away.

**Figure 5:**
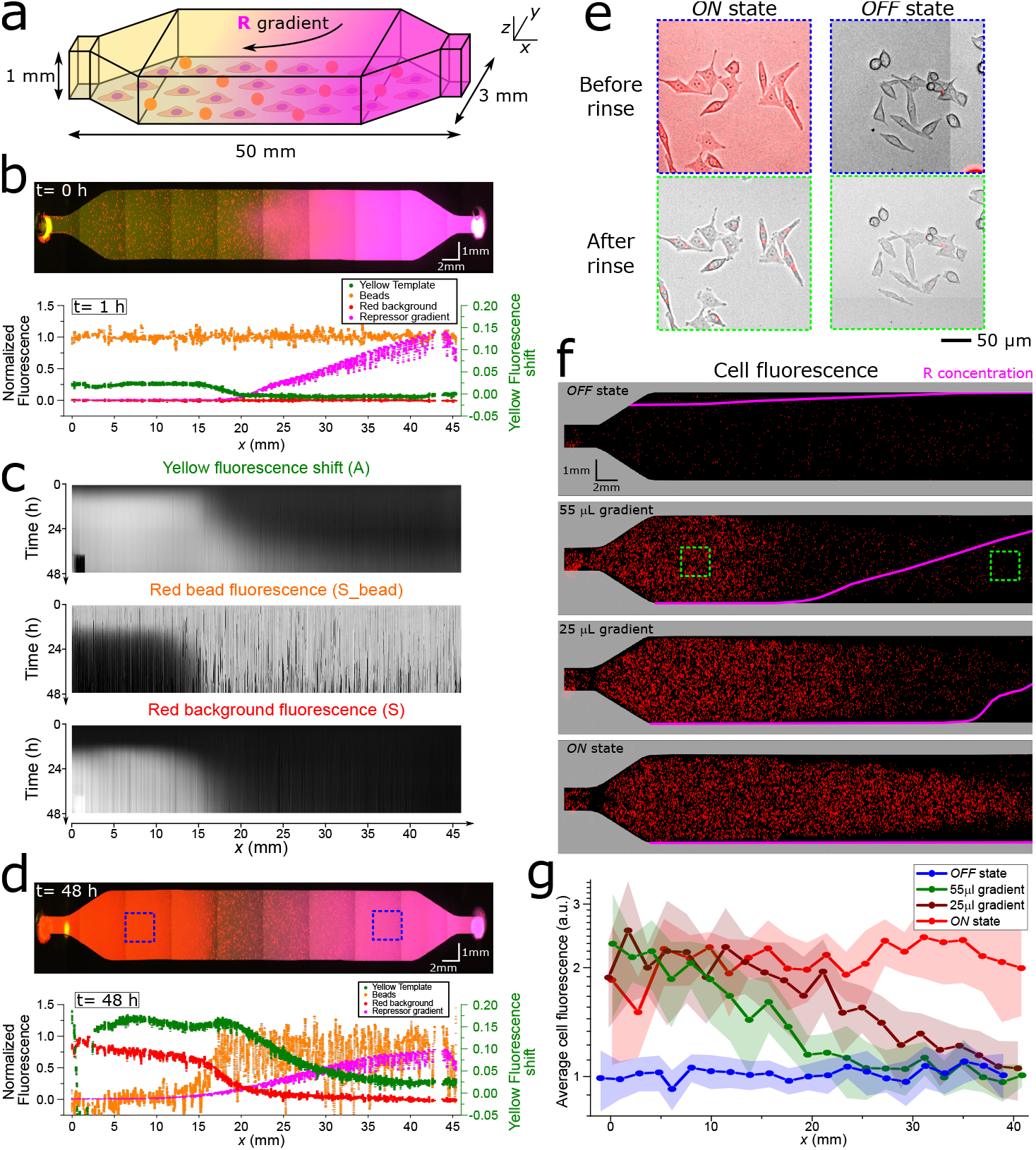
The reactive medium processes extracellular positional information and transfers it to the cells. a) Scheme (not to scale) of the millifluidic channel where cells, conversion beads and the medium containing the internalization switch (in yellow) are homogeneously distributed and the repressor (in magenta) forms a gradient along the *x* axis. b) Three-colour fluorescence image of the channel at *t* = 0 h and corrected fluorescence profiles at *t* = 1 h. c) Kymographs of the fluorescence profiles associated to species **A** (top), **S** on the beads (middle) and **S** in solution (bottom) d) Fluorescence image of the channel and corresponding profiles at *t* = 48 h. e) Composite bright-field/red fluorescence images of the cells at *t* = 48 h in regions where the switch is ON and OFF, before and after rinsing. The corresponding regions are highlighted in panels d and f with dashed blue and green rectangles, respectively. f) Red fluorescence images of the rinsed cells for different initial gradients of **R** represented as magenta lines. The grey zones correspond to the position of the channel walls before rinsing. g) Profiles of the average fluorescence per cell corresponding to the four gradients of **R** in panel f. 10 cells were measured for each point. Shades correspond to the standard deviation. To help visualization, beads profiles in panels b and d have been smoothed by an adjacent averaging of 50 data points. Panels b-e correspond to experiments with a 55 *μ*L **R** gradient. Conditions: [A]_0_ = 20 nM, [T]_0_ = 200 nM and maximal [R]= 2 *μ*M.

Figure 5c shows fluorescence kymographs related to the concentration of **A** in the medium (yellow) and **S** on the beads (red bead) and in the medium (red background) during this experiment. The autocatalytic production of **A** started on the left side of the channel (*x* < 17 mm), where the concentration of **R** was zero, and generated a steady-state two-band pattern between t = 10 h and 23 h. At t = 10 h, when **A** reached its steady-state, it triggered the release of **S** from the beads into the medium for cellular internalization, which also formed a two-band pattern. After 23 h, **A** propagated in the form of a reaction-diffusion front with a velocity of 4.3 *μ*m/min, while the pattern of **S** on the beads was static and its release into the medium showed minor propagation.

In the absence of cells, the spatio-temporal behaviour of the internalization switch was slightly different (Figure S17). In this case, and depending on the initial shape of the gradient, the two-band pattern of **A** either stopped or a faint front directly occurred, while the front of released **S** did not stop but propagated at constant velocity. These differences may be due to the degradation of the DNA species by the cells, an effect that should be more important in spatial than in temporal experiments because, for geometrical reasons, the cell volume density was 4-fold higher in the former. Indeed, the front of **A** that arises at long times in the presence of cells could be attributed to the onset of diffusion-independent autocatalysis due to a degradation of **R** by the cells. In addition, the front of **S** could be halted by cell internalization combined to the non-linear conversion of **A** into **S** (Figure S10). This could further explain why at *t* = 48 h, in the presence of cells, the profile of **S** in solution is sharper than the profile of **A** (Figure 5d) – 5 and 15 mm width, respectively.

Despite these differences, the spatial version of the internalization switch works as designed in the presence of cells: it creates a two-band pattern of **S**. In addition, Figure 5e shows that, after 48 h, the majority of cells become red on the left side of the pattern, while the inverse is true on the right side. To test that cell modification was truly controlled by positional information, we generated four different gradients of **R**. Figure 5f and g display the images and corresponding profiles along the channel axis of the red fluorescence inside the cells for different initial conditions of **R** (Figure S18 shows the kymographs for the other species). In agreement with the temporal experiments above, in the presence of respectively high and zero concentration of **R**, the cells appear dark or red along the whole channel. In contrast, the presence of a gradient of **R** made cells become red only in the zone where **S** was released. Importantly, the position of the gradient of **R** determined the width of the band of released S, and hence of that of red cells. The border between red and dark cells was, however, twice wider than that of red fluorescence in solution at final time, since cells internalize **S** over time (Figure 4). Taken together, these results demonstrate that it is possible to engineer an out-of-equilibrium reactive medium that takes as input an extracellular source of positional information that cannot be sensed by the cells, and that processes and translates it into a chemical signal that is readily internalized by living cells.

## Conclusions

In this work, we have developed an extracellular medium that sustains out-of-equilibrium reaction networks with feedbacks. We have shown that such networks change the composition of living cells with high control both in time and space. To do so, we have found common conditions in which an extracellular DNA-enzyme program was functional in the presence of a viable culture of human cells. With these tools we have engineered an reactive extracellular medium that controls the internalization of a DNA strand inside living cells. The timing of internalization can be set either externally by adding a reagent or internally by tuning the initial composition of the medium and taking advantage of a clock reaction. In addition, we have demonstrated that reactive extracellular media may create concentration patterns that spatially control the internalization of DNA by cells, mimicking the transfer of positional information at play during early embryo development. ^27,34,46^ Importantly, these patterns were generated autonomously by non-equilibrium chemical reactions without hydrodynamic flow, in contrast with standard methods relying on microfluidics, ^47^ where flow introduces shear stress and washes out nutrients and signaling molecules.

Our method stands out for its simplicity. Non-equilibrium DNA programs could be run in the presence of living cells without introducing any major modification other than a variation in the composition of a classical growth medium. In particular, no chemical modification of the DNA program, no genetic modification of the cells, nor special apparatus other than standard cell culture well-plates were needed to run the autocatalytic network and the bistable switch. Only the more complex internalization program needed the addition of hydrogel beads to physically separate the cells from the output DNA strand. Remarkably, the program worked at 37 °C and without human intervention for at least 48 h. The presence of the enzymes that continuously synthesize fresh DNA may account for the robustness of the method, although complementary approaches based on DNA strand displacement reactions may improve the programmability of the reactive medium. ^48^ In addition, the DNA programs were fast compared with cell division (3 to 7-fold), and remained active for two days, which could be possibly extended up to a week. ^35^ Furthermore, the wide array of reactivities offered by oligonucleotides, thanks to chemical modification or sequence selection, anticipates that the reactive medium developed here could receive and release a variety of inputs and outputs that would further control cellular behaviour, either genetically^49^ or via aptamers.^10^ Finally, the out-of-equilibrium dynamics of the medium offer an advantage to build devices responsive to a change in the cellular state.

With these properties, one foresees that DNA-based reactive extracellular media could be used for real-time *in vitro* detection of fast evolving or long-lasting biomarkers in living cells. ^50^ In addition, extracellular self-organizing DNA concentration patterns could provide novel methods to engineer tissues, with differentiation induced over space and time in a controlled manner. Ultimately, reactive media could be integrated into smart bandages^8^ or into engineered living materials^51^ but also function as a detector,^50^ as an actuator, or both, while benefiting of the computation capabilities of DNA.

## Methods

All DNA strands were designed heuristically and with the help of Nupack^52^ and purchased from Integrated DNA Technologies, Inc (U.S.) or Biomers (Germany). The Bst DNA polymerase large fragment and the Nb.BsmI nicking enzymes were purchased from New England Biolabs, while the *Thermus thermophilus* RecJ exonuclease was produced in-house as described. ^53^

The DNA buffer was used as described previously. ^34^ The reference cell-buffer contained DMEM supplemented with 1% Penicillin-Streptomycin.The cell-DNA buffer contained all the components of DMEM at 2-fold dilution, 1% Penicillin-Streptomycin and all the constituents of the DNA buffer at their standard concentration, except for (NH_4_)_2_SO_4_, that was suppressed, and NaCl and dithiothreitol (DTT), whose concentrations were lowered, to reduce toxicity (see Table S1). Unless otherwise stated, all experiments were performed at 37 °C. DNA sequences and further experimental procedures are provided in the Supplementary Information.

### Monitoring of PEN DNA reactions

Standard enzyme concentrations for PEN reactions were 8 U/mL polymerase, 100 U/mL nickase, 31.25 nM exonuclease, and 0.4 mM dNTPs (New England Biolabs). When cell culture conditions were not needed (Figure 1b), the dynamics of PEN reactions were recorded in 20 *μ*L solutions inside 150 *μ*L qPCR tubes using a Qiagen Rotor-Gene qPCR machine or a CFX96 Touch Real-Time PCR Detection System (Bio-Rad). For biological relevance, homogeneous experiments with cells were performed in well plates (see below) and their fluorescence monitored by microscopy. To calculate the fluorescence shift, the raw fluorescence intensity was normalized by an early time point (*t* = 5 min), and subtracted from 1, as done previously.^34^ The onset amplification time, **τ**, was defined as the time point at which 50% of the fluorescence of the steady state of the *ON* state was reached.

Polyacrylamide denaturing gel electrophoresis at 20% was run for 2 h at 200 V in 0.5X TBE buffer, stained with 1000x Sybr Gold (ThermoFisher: S11494) for 10 min, and imaged using a Gel Doc^™^ EZ Gel Imager (Bio-Rad). Note that we use species **A**^1^ because upon the hydrolysis of a phosphodiester bond during the nicking event, the phosphate group remains in the 5’ of the second trigger, since if the phosphate group remained on the 3’ of the first trigger no autocatalytic behaviour would be attainable.

### Spatial experiment devices

The 50 mm long, 3 mm wide and 1 mm high millifluidic channels were made out of poly(dimethylsiloxane) (PDMS, RTV 615). Firstly, a milling machine is used to create a reusable polyvinyl chloride (PVC) mold (Figure S19). PDMS (prepolymer/curing agent ratio (w/w) of 10: 1) is subsequently poured into the mold, degassed in a vacuum chamber to remove the bubbles, and cured at 65°C for 1.5 hours. Once done, the PDMS layer is removed from the mold, access holes are punched on both ends of the channels, and the PDMS is gently pressed on a glass slide, both surfaces being previously plasma treated.

### Cell culture handling and experiments

Human cervix epitheloid carcinoma cells (HeLa cell line) were grown at 37 °C and 5% CO_2_. The cells were firstly grown in two sequential steps (10% and 5%) of FBS (Dominique Dutscher: S1810-500) before reducing the FBS down to 2.5%, to avoid drastic shock upon removal of FBS. The adherent culture was maintained in 2.5% FBS cell-buffer medium until reaching 80-90% confluence, where cells were trypsinized with trypsin-EDTA (PAN Biotech: P10-019100) and diluted into fresh 2.5% FBS cell-buffer medium. Experiments carried out in 384 cell well plates (ThermoFisher: 142762) were seeded with 1600 cells in 50 *μ*L of medium per well. For spatial experiments, cells were seeded within the millifluidic channels using a 200 *μ*L pipette tip at 1.1×10^5^ cells/ml. In both cases, the cells were allowed to adhere to the surface for 24 h before further experimental handling.

For cell viability experiments, the medium was removed from the well and replaced with 50 *μ*L of the buffer under test, and left for the stated number of days. For DMSO controls, 10 *μ*L of DMSO (Panreac: A3672) was directly added from stock solution. For trypan blue viability assays, 2 *μ*L of trypan blue 0.4% solution (Gibco: 15250061) was added and gently mixed. After 5 min in the incubator, the wells were rinsed twice with 1X PBS, always leaving some solution in the well to assure that no cells were removed during rinsing. Cells were then imaged in bright-field with a 2.5X 5324.8 *μm*^2^ and a 20X 644.84 *μm*^2^ objective.

For quantification of cells by fluorescent-activated cell sorting (FACS), the wells were gently rinsed with 50 *μ*L 1X DMEM and incubated 8 minutes with 50 *μ*L trypsin (stock solution) before inactivation with 50 *μ*L of DMEM supplemented with 10% FBS. The cell solution was mixed with 250 *μ*L FACSFlow (Fisherscientific: 12756528) and 0.5 *μ*L of propidium iodide (20 mM, ThermoFisher: L7012). Heat-treated cells were left at 65 °C for 10 minutes. A Becton-Dickinson flow cytometer (FACSCalibur) equipped with a 488 nm argon ion laser was used to excite propidium iodide, and record the emissions in the fluorescence channel FL-2 (band pass 585/42 nm). Cells were quantified for 3 min at a flow of 60 *μ*L/min. Fluorescent measurements were treated and analysed with a home-made Matlab (The Mathworks) routine. Note: due to the presence of a reducing agent (DTT), standard MTT viability test could not be performed.

Concerning experiments with cells in the presence of the reactive medium, for 384 well plate experiments the cell culture medium was removed and replaced with 50 *μ*L of the reactive medium. For spatial experiments, the reactive medium was injected twice gently to remove the cell culture medium without detaching the cells. The channel inlets and outlets were closed with greases, and a coverslide was pressed onto the grease for a better sealing limiting evaporation. In the case of inhomogeneous initial conditions (gradient of **R**), two identical solutions are prepared where only one is supplemented with **R**. The channel is first filled with the solution without **R** and left 5 minutes to allow the beads to sediment to the surface. Then, a precise volume of the solution containing **R** is injected. A gentle back and forth pipetting is performed 5 times to generate a gradient of **R** along the channel, while all other components remain homogeneously distributed. Figure S19 shows, with the use of methylene blue dye, the visual appearance of the gradients obtained using this setup. For visualization of the **R** gradients, **R**^1^ was used at 10% of the desired final **R** concentration.

### Particle loading

Conversion beads were constituted of porous streptavidin-conjugated sepharose hydrogel with 34 *μ*m average particle size (GE healthcare: GE17-5113-01) functionalized with biotinylated DNA constructs. Firstly, we annealed 290 pmoles of double-biotinylated **C_1_** with 247 pmoles of **S** in a 115.7 *μ*L 5 mM MgSO_4_ solution with a temperature ramp of 1 °C every 10 s, from 90 °C to 20 °C. Secondly, we rinsed twice 5 *μ*L of Sepharose solution (~10000 beads/μL) with washing buffer (10 mM Tris-HCl, pH 7.5, 2 M NaCl, 1 mM EDTA, 0.2% Tween 20), followed by a third wash with deionized water and decantation. The DNA annealed solution was mixed with the rinsed beads and immediately agitated for 10 seconds, to further incubate it at 40 °C for 30 min, with agitation to avoid particle sedimentation. Finally, the functionalized beads were rinsed three times with a 5 mM MgSO**4** solution to remove unbound DNA, and the final volume was adjusted to 100 *μ*L. Functionalized beads were used at 0.62% and 1.67% for the 384 well plate and the spatial experiments respectively, which accounted for low surface coverage (Figure S9).

### Microscopy

PEN DNA reactions (with and without cells) were monitored using a Zeiss Axio Observer Z1 fully automated epifluorescence microscope equipped with a ZEISS Colibri 7 LED light, YFP and RFP filter sets, and a Hamamatsu ORCA-Flash4.0V3 inside a Zeiss incubation system to regulate temperature at 37 °C, in the presence of high humidity and at 5% CO_2_. Note: we observed up to a 75% reduction in *τ* for 384 well plates (Figure 2a) than that for qPCR machines (Figure 1b), which we mainly attribute to a difference in the materials of the two containers used (polystyrene wells and polypropylene tubes respectively) and to a lower robustness of the microscope to maintain a constant temperature compared to a thermocycler. Multi-color fluorescence and bright-field images were recorded every 10 min (Figure 2), 35 min (Figure 4) or 1 h (Figure 3) with a 2.5X objective, and every 1 h with a 72-by-1 image array using a 10X 648.1 *μ*m^2^ objective for Figure 5. For spatial experiments, prior and following temporal dynamics, the entire millifluidic channel was imaged with a 2.5X objective.

To detect cellular internalization, cells in 384 well plates were rinsed twice with 60 *μ*L of DMEM (with a 5 min incubation at 37 °C between rinses) and the entire well was imaged with a 4-by-4 image array using a 20X objective. For cells in the millifluidic channel, the PDMS device was removed and each channel was rinsed twice with 1 mL of DMEM, to later be imaged with a 40-by-4 image array using a 10X objective (Figure 5f), and a 72-by-1 image array using a 20X objective (Figure 5g).

### Image and data treatment

The raw microscopy images were treated with ImageJ / Fiji (NIH) and Matlab. To follow the dynamics of the PEN reactions, time-lapse images were imported into a stack for each well. Three threshold values were defined to select the template fluorescence (yellow channel), the background fluorescence and the bead fluorescence (red channel). In the case of the beads, pixels were only selected if they stayed within the selection criteria for more than half the duration of the experiment, to avoid disturbances caused by the movement of the beads. Each thresholded image was averaged along the *y* axis to create an intensity profile at each time point. These profiles were stacked over time into a kymograph, which was then normalized by the profile of an initial time point to correct from inhomogeneous illumination. The kymographs were then averaged over space to obtain the profiles of intensity versus time. To correct from time-dependent artefacts introduced by the microscope when performing experiments in 384 well plates, the profiles of each well were normalized with a negative control (absence of enzymes, *n* = 2). In spatial experiments, the inhomogeneous illumination in the repressor (**R**) gradients was corrected by dividing each frame of the channel by a frame in the absence of **R** (left most side of the channel). Furthermore, to adjust all the *x* positions of the kymographs to the same spatial position, the kymographs corresponding to the template fluorescence, the background fluorescence and the **R** gradient were re-plotted with only the same *x* position as the kymograph of the beads. Fluorescence shifts were calculated as described above.

For the characterization of cellular fluorescence of experiments performed in 384 well plates, and prior to combining images into the 4-by-4 image array, the background of red fluorescence images was subtracted using ImageJ ‘subtract background’ routine with a 2000 pixel rolling ball radius and sliding paraboloid parameters. Subsequently, the mean background fluorescence was removed from the image to have a homogenous background within the image array. The contour of each cell was manually selected in the bright-field image and used to extract the average red fluorescence of each cell. Selection of cells was restricted to the middle of the wells to avoid optical artefacts. For spatial experiments, due to higher efficacy of the rinsing, the average red fluorescence of each cell could be directly quantified from the image.

## Supporting information

The Supporting Information contains further experimental methods, a discussion of the buffer screening process and details on the composition, DNA sequences, 19 supporting figures and 1 supporting video.

## Acknowledgements

We thank Stéphanie Bonneau and Ramón Eritja for supplying the cells, Clara Berenguer-Escuder for advice with cell culture, and Matthieu Morel, Yannick Rondelez and Guillaume Ginés for insightful discussions. We also thank the financial support from the European Research Council (ERC) under the European’s Union Horizon 2020 program (grant no. 770940, A.E.-T.), by the Ville de Paris Emergences program (Morphoart, A.E.-T.), by a Marie Sklodowska-Curie fellowship (grant no. 795580, M.vdH.) from the European Union’s Horizon 2020 program, and by a PRESTIGE grant (grant no. 609102, M.vdH.) from the European Union’s Seventh Framework Programme.

## Author contributions

All authors designed research, discussed the results and wrote the paper. M.vdH. designed and performed the experiments and analysed the data.

## TOC figure

**Figure 6:**
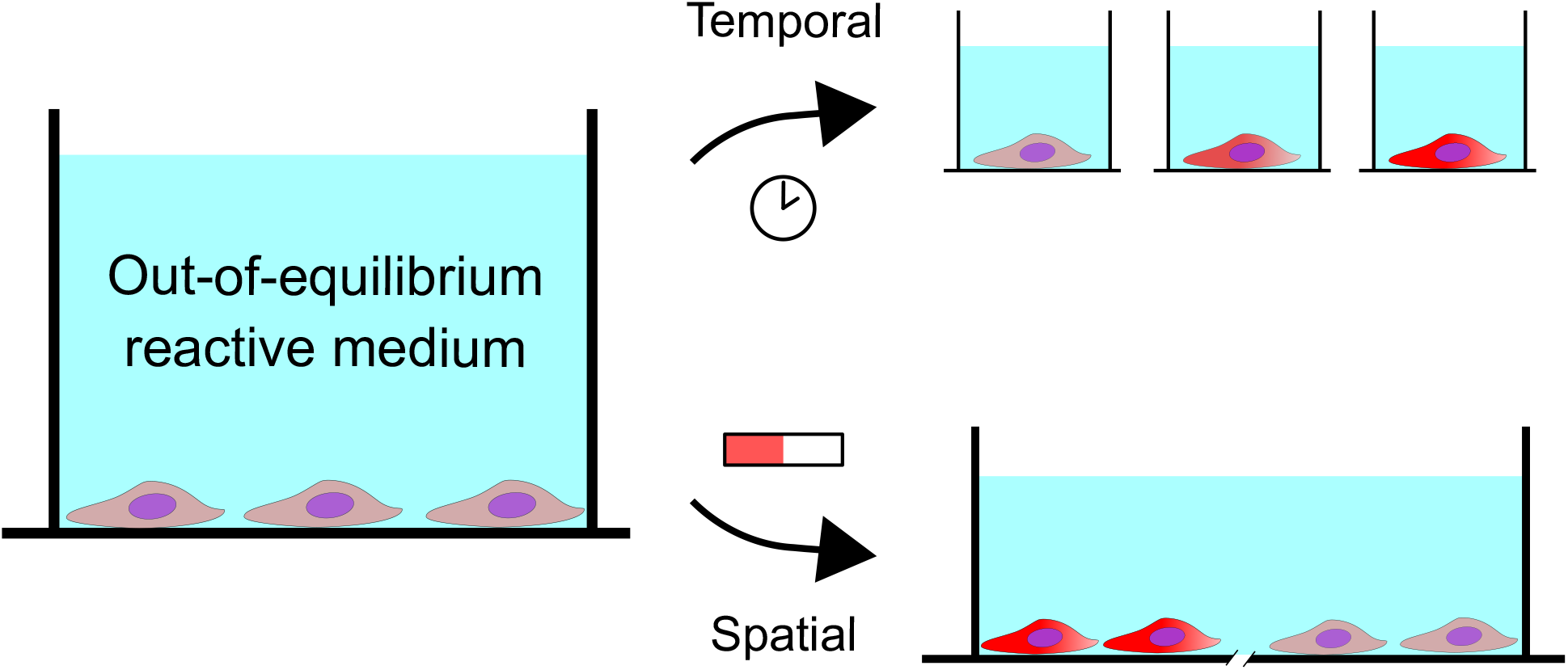
For Table of Contents Only

## References

(1) Cameron, D. E.; Bashor, C. J.; Collins, J. J. A brief history of synthetic biology. Nat Rev Micro 2014, 12, 381–390.

(2) Kelly, C. L.; Liu, Z.; Yoshihara, A.; Jenkinson, S. F.; Wormald, M. R.; Otero, J.; Estévez, A.; Kato, A.; Marqvorsen, M. H. S.; Fleet, G. W. J.; Estévez, R. J.; Izumori, K.; Heap, J. T. Synthetic Chemical Inducers and Genetic Decoupling Enable Orthogonal Control of the rhaBAD Promoter. ACS Synthetic Biology 2016, 5, 1136–1145.

(3) Wan, X.; Marsafari, M.; Xu, P. Engineering metabolite-responsive transcriptional factors to sense small molecules in eukaryotes: current state and perspectives. Microbial Cell Factories 2019, 18, 61.

(4) Shimizu-Sato, S.; Huq, E.; Tepperman, J. M.; Quail, P. H. A light-switchable gene promoter system. Nature Biotechnology 2002, 20, 1041–1044.

(5) Toda, S.; Blauch, L. R.; Tang, S. K. Y.; Morsut, L.; Lim, W. A. Programming self-organizing multicellular structures with synthetic cell-cell signaling. Science 2018, 361, eaat0271.

(6) Greber, D.; Fussenegger, M. An engineered mammalian band-pass network. Nucleic Acids Research 2010, 38, e174–e174.

(7) Johnson, M. B.; March, A. R.; Morsut, L. Engineering multicellular systems: using synthetic biology to control tissue self-organization. Current opinion in biomedical engineering 2017, 4, 163–173, 29308442 [pmid] PMC5749256 [pmcid].

(8) Derakhshandeh, H.; Kashaf, S. S.; Aghabaglou, F.; Ghanavati, I. O.; Tamayol, A. Smart Bandages: The Future of Wound Care. Trends in Biotechnology 2018, 36, 1259–1274.

(9) Acar, M.; Mettetal, J. T.; Van Oudenaarden, A. Stochastic switching as a survival strategy in fluctuating environments. Nature Genetics 2008, 40, 471–475.

(10) Chen, Y. J.; Groves, B.; Muscat, R. A.; Seelig, G. DNA nanotechnology from the test tube to the cell. Nature Nanotechnology 2015, 10, 748–760.

(11) Douglas, S. M.; Bachelet, I.; Church, G. M. A logic-gated nanorobot for targeted transport of molecular payloads. Science 2012, 335, 831–834.

(12) Li, S. et al. A DNA nanorobot functions as a cancer therapeutic in response to a molecular trigger in vivo. Nature Biotechnology 2018, 36, 258–264.

(13) Freeman, R.; Stephanopoulos, N.; Álvarez, Z.; Lewis, J. A.; Sur, S.; Serrano, C. M.; Boekhoven, J.; Lee, S. S.; Stupp, S. I. Instructing cells with programmable peptide DNA hybrids. Nature Communications 2017, 8, 15982.

(14) Bhatia, D.; Arumugam, S.; Nasilowski, M.; Joshi, H.; Wunder, C.; Chambon, V.; Prakash, V.; Grazon, C.; Nadal, B.; Maiti, P. K.; Johannes, L.; Dubertret, B.; Krishnan, Y. Quantum dot-loaded monofunctionalized DNA icosahedra for single-particle tracking of endocytic pathways. Nature Nanotechnology 2016, 11, 1112–1119.

(15) Rudchenko, M.; Taylor, S.; Pallavi, P.; Dechkovskaia, A.; Khan, S.; Butler, V. P.; Rudchenko, S.; Stojanovic, M. N. Autonomous molecular cascades for evaluation of cell surfaces. Nature Nanotechnology 2013, 8, 580–586.

(16) You, M.; Zhu, G.; Chen, T.; Donovan, M. J.; Tan, W. Programmable and multiparameter DNA-based logic platform for cancer recognition and targeted therapy. Journal of the American Chemical Society 2015, 137, 667–674.

(17) Song, T.; Shah, S.; Bui, H.; Garg, S.; Eshra, A.; Fu, D.; Yang, M.; Mokhtar, R.; Reif, J. Programming DNA-Based Biomolecular Reaction Networks on Cancer Cell Membranes. Journal of the American Chemical Society 2019, 141, 16539–16543.

(18) Rinaudo, K.; Bleris, L.; Maddamsetti, R.; Subramanian, S.; Weiss, R.; Benenson, Y. A universal RNAi-based logic evaluator that operates in mammalian cells. Nature Biotechnology 2007, 25, 795–801.

(19) Chatterjee, G.; Chen, Y. J.; Seelig, G. Nucleic Acid Strand Displacement with Synthetic mRNA Inputs in Living Mammalian Cells. ACS Synthetic Biology 2018, 7, 2737–2741.

(20) Lentini, R.; Santero, S. P.; Chizzolini, F.; Cecchi, D.; Fontana, J.; Marchioretto, M.; Del Bianco, C.; Terrell, J. L.; Spencer, A. C.; Martini, L.; Forlin, M.; Assfalg, M.; Serra, M. D.; Bentley, W. E.; Mansy, S. S. Integrating artificial with natural cells to translate chemical messages that direct E. coli behaviour. Nature Communications 2014, 5, 4012.

(21) Schwarz-Schilling, M.; Aufinger, L.; Mückl, A.; Simmel, F. C. Chemical communication between bacteria and cell-free gene expression systems within linear chains of emulsion droplets. Integrative Biology (United Kingdom) 2016, 8, 564–570.

(22) Lentini, R.; Martín, N. Y.; Forlin, M.; Belmonte, L.; Fontana, J.; Cornella, M.; Martini, L.; Tamburini, S.; Bentley, W. E.; Jousson, O.; Mansy, S. S. Two-Way Chemical Communication between Artificial and Natural Cells. ACS Central Science 2017, 3, 117–123.

(23) Toparlak, O. D.; Zasso, J.; Bridi, S.; Serra, M. D.; Macchi, P.; Conti, L.; Baudet, M.-L.; Mansy, S. S. Artificial cells drive neural differentiation. Science Advances 2020, 6.

(24) Niederholtmeyer, H.; Stepanova, V.; Maerkl, S. J. Implementation of cell-free biological networks at steady state. Proceedings of the National Academy of Sciences 2013,

(25) Karzbrun, E.; Tayar, A. M.; Noireaux, V.; Bar-Ziv, R. H. Programmable on-chip DNA compartments as artificial cells. Science 2014, 345, 829–832.

(26) Briscoe, J.; Small, S. Morphogen rules: design principles of gradient-mediated embryo patterning. Development 2015, 142, 3996–4009.

(27) Urtel, G.; Van Der Hofstadt, M.; Galas, J. C.; Estevez-Torres, A. REXPAR: An Isothermal Amplification Scheme That Is Robust to Autocatalytic Parasites. Biochemistry 2019, 58, 2675–2681.

(28) Montagne, K.; Plasson, R.; Sakai, Y.; Fujii, T.; Rondelez, Y. Programming an in vitro DNA oscillator using a molecular networking strategy. Mol Syst Biol 2011, 7, 466, 10.1038/msb.2010.120.

(29) Van Der Hofstadt, M.; Gines, G.; Galas, J.-C.; Estevez-Torres, A. In DNA- and RNA-Based Computing Systems; Evgeny, K., Ed.; John Wiley & Sons, in press.

(30) Fujii, T.; Rondelez, Y. Predator-Prey Molecular Ecosystems. ACS Nano 2013, 7, 27–34.

(31) Padirac, A.; Fujii, T.; Rondelez, Y. Quencher-free multiplexed monitoring of DNA reaction circuits. Nucleic Acids Research 2012, 40, e118–e118.

(32) Montagne, K.; Gines, G.; Fujii, T.; Rondelez, Y. Boosting functionality of synthetic DNA circuits with tailored deactivation. Nature Communications 2016, 7, 13474.

(33) Padirac, A.; Fujii, T.; Estevez-Torres, A.; Rondelez, Y. Spatial Waves in Synthetic Biochemical Networks. Journal of the American Chemical Society 2013, 135, 14586–14592, doi:10.1021/ja403584p.

(34) Zadorin, A. S.; Rondelez, Y.; Gines, G.; Dilhas, V.; Urtel, G.; Zambrano, A.; Galas, J. C.; Estevez-Torres, A. Synthesis and materialization of a reaction-diffusion French flag pattern. Nature Chemistry 2017, 9, 990–996.

(35) Urtel, G.; Estevez-Torres, A.; Galas, J.-C. DNA-based long-lived reaction–diffusion patterning in a host hydrogel. Soft Matter 2019, 15, 9343–9351.

(36) Baccouche, A.; Montagne, K.; Padirac, A.; Fujii, T.; Rondelez, Y. Dynamic DNA-toolbox reaction circuits: A walkthrough. Methods 2014, 67, 234–249.

(37) Xiang, X.-Y.; Yang, X.-C.; Su, J.; Kang, J.-S.; Wu, Y.; Xue, Y.-N.; Dong, Y.-T.; Sun, L.-K. Inhibition of autophagic flux by ROS promotes apoptosis during DTT-induced ER/oxidative stress in HeLa cells. Oncology Reports 2016, 35, 3471–3479.

(38) Epstein, I.; Pojman, J. A. An introduction to nonlinear chemical reactions; Oxford University Press: New York, 1998.

(39) Zhao, Q.; Matson, S.; Herrera, C. J.; Fisher, E.; Yu, H.; Krieg, A. M. Comparison of Cellular Binding and Uptake of Antisense Phosphodiester, Phosphorothioate, and Mixed Phosphorothioate and Methylphosphonate Oligonucleotides. Antisense Research and Development 1993, 3, 53–66.

(40) Gines, G.; Zadorin, A. S.; Galas, J. C.; Fujii, T.; Estevez-Torres, A.; Rondelez, Y. Microscopic agents programmed by DNA circuits. Nature Nanotechnology 2017, 12, 351–359.

(41) Zhang, D. Y.; Seelig, G. Dynamic DNA nanotechnology using strand-displacement reactions. Nat Chem 2011, 3, 103–113, 10.1038/nchem.957.

(42) Nödling, A. R.; Mills, E. M.; Li, X.; Cardella, D.; Sayers, E. J.; Wu, S.-H.; Jones, A. T.; Luk, L. Y. P.; Tsai, Y.-H. Cyanine dye mediated mitochondrial targeting enhances the anti-cancer activity of small-molecule cargoes. Chemical Communications 2020,

(43) Zadorin, A. S.; Rondelez, Y.; Galas, J.-C.; Estevez-Torres, A. Synthesis of Programmable Reaction-Diffusion Fronts Using DNA Catalyzers. Phys. Rev. Lett. 2015, 114, 068301.

(44) Wolpert, L. Positional information and the spatial pattern of cellular differentiation. J. Theor. Biol. 1969, 25, 1–47.

(45) Akam, M. Hox and HOM: Homologous gene clusters in insects and vertebrates. Cell 1989, 57, 347–349.

(46) Senoussi, A.; Vyborna, Y.; Berthoumieux, H.; Galas, J.-C.; Estevez-Torres, A. In Out-of-equilibrium supramolecular systems and materials; Giuseppone, N., Walther, A., Eds.; Wiley-VCH, in press.

(47) Kim, S.; Kim, H. J.; Jeon, N. L. Biological applications of microfluidic gradient devices. Integrative Biology 2010, 2, 584–603.

(48) Fern, J.; Schulman, R. Design and Characterization of DNA Strand-Displacement Circuits in Serum-Supplemented Cell Medium. ACS Synthetic Biology 2017, 6, 1774–1783.

(49) Lundin, K. E.; Gissberg, O.; Smith, C. E. Oligonucleotide Therapies: The Past and the Present. Human Gene Therapy 2015, 26, 475–485.

(50) Gines, G.; Menezes, R.; Nara, K.; Kirstetter, A.-S.; Taly, V.; Rondelez, Y. Isothermal digital detection of microRNAs using background-free molecular circuit. Science Advances 2020, 6.

(51) Guo, S. et al. Engineered Living Materials Based on Adhesin-Mediated Trapping of Programmable Cells. ACS Synthetic Biology 2020, 9, 475–485.

(52) Zadeh, J. N.; Steenberg, C. D.; Bois, J. S.; Wolfe, B. R.; Pierce, M. B.; Khan, A. R.; Dirks, R. M.; Pierce, N. A. NUPACK: Analysis and design of nucleic acid systems. Journal of Computational Chemistry 2011, 32, 170–173.

(53) Wakamatsu, T.; Kitamura, Y.; Kotera, Y.; Nakagawa, N.; Kuramitsu, S.; Masui, R. Structure of RecJ exonuclease defines its specificity for single-stranded DNA. J Biol Chem 2010, 285, 9762–9.

