## supplementary Information for "Spatio-temporal patterning of living cells with extracellular DNA programs"

### Contents


1 Methods 3

2 Buffer screening 5

3 Tables S1 to S2 11

4 Figures S5 to S14 14

5 Video S1 29

References 30

### 1 Methods

#### 1.1 Determining the cytotoxicity of the PEN DNA toolbox components

Each component used in standard PEN reactions[1] was individually tested to know its relative cytotoxicity on living HeLa cells. 3200 cells were cultured in 96 cell well plates with 2.5% FBS for 24h, allowing cells sufficient time to adhere to the surface. The medium was then replaced with 100 $\mu$L of medium containing 1% Penicillin-Streptomycin, Dulbecco's modified Eagle's medium (DMEM) and the compound of interest at the stated concentration. Viability was determined by the visual appearance of cells after the specified incubation times. Bright-field images (581x486 $\mu m^2$) were taken with an Olympus CK2 Inverted Microscope, 10X objective and a Point Grey BFS-U3-51S5M-C camera. Images represent the general appearance of cells at each tested condition (per duplicate).

#### 1.2 Determining reaction rates of individual enzymes

The reaction rates for each enzyme were measured in different buffers and temperatures by following the fluorescence signal over time of a reference substrate in the presence of the enzyme. For the BsmI nicking enzyme, the reference substrate was a DNA molecular beacon whose stem carried the nicking enzyme recognition site ($ref_1$). The nicking event caused the release of a short fluorescent oligonucleotide, and hence an increase in the fluorescence along the reaction. For the exonuclease, the reference substrate was a 16-mer oligonucleotide ($ref_2$) at 1 $\mu$M in the presence of a fluorescent DNA intercalator (EvaGreen, Biotium). The degradation of this oligonucleotide by the exonuclease causes a reduction in fluorescence over time.

The fluorescence signals were measured from 20 $\mu$L solutions using a Bio-rad CFX qPCR: it increased (or decreased) linearly until reaching a plateau. Raw fluorescence signals were normalized between 0 and 1 and converted to DNA concentrations by multiplying them by the substrate concentration. These treated data were fitted by a linear function and the slope provided the rates displayed in Figure S6.

#### 1.3 Determining trigger concentrations at steady states.

The steady-state trigger concentration ($[\mathrm{A}]_{ss}$) was measured at 37 °C in the Rotor-Gene qPCR machine for both the **A** and the $\mathbf{A}_1$ amplification sequences as previously described.[2] Firstly, 20 $\mu$L of exponential amplification was left until it reached steady state (900 and 1200 min in this case) with 50nM of template and 0.5nM of initial trigger concentrations. Secondly, an extract from this solution (1% and 5% in this case) was introduced into a fresh exponential reaction. Simultaneously, a calibration curve was done with a range of the initial trigger concentration, to extract the autocatalytic rate from the amplification onset times. Using the autocatalytic rate, the predicted initial trigger concentration of the condition under test can be extracted from their amplification onset times. The initial trigger concentration is then multiplied by 100 or 20 (for 1% and 5% extracts, respectively) to determine the trigger concentrations at steady states in Figure S5.

### 2 Buffer screening

Both living cells and the PEN DNA toolbox rely on biochemical reactions to sustain, but each has its own buffer composition. Living cells require a rich nutritious medium, while that of the PEN DNA toolbox is relatively simple and admits variations.[3] Due to a higher flexibility and robustness of PEN reactions,[4] we decided in prioritizing cellular viability over PEN reactions' efficiency when attaining a buffer compatible to cell growth and PEN reactions.

Firstly, we screened the standard PEN DNA toolbox for cytotoxic components, with interest in obtaining a biocompatible enzyme-based DNA program without significant cell death for at least 3 days. To do so, we broke down the DNA buffer into its components to assess each component individually, in addition of testing the cytotoxicity of DNA and enzymes (Figure S1 and Figure S2). After 22 h, we already observed a high mortality of HeLa cells in the presence of the PP buffer (a home made buffer). The PP buffer cannot be entirely removed, since it contains the essential compounds for the PEN reactions; the energy source (dNTPs), and $MgSO_4$, which is essential for DNA dynamics.[5] For this reason, the PP buffer was further broken down, and each constituent was tested individually at their working concentration (Figure S1 middle panel). While the essential components for the PEN reactions didn't present cell toxicity (dNTPS and $MgSO_4$), after 47 h the cytotoxic components of the PP buffer were evident. The addition of sodium chloride, or of Thermopol buffer (home made variation to the commercial one) presented significant cell death. Since the cell culture medium (DMEM) contains 120 mM of sodium chloride, we deducted that the further addition of 50 mM could increase drastically osmolarity, and hence cause cellular death. As DMEM contains more than double what is needed for the PEN reactions, we decided that the further addition of sodium chloride was not needed. For the other cytotoxic component of the PP buffer, the Thermopol buffer, we repeated the procedure and broke it down into its constituents, and further tested them individually (Figure S1 bottom panel). Here, we found that ammonium sulphate was the cytotoxic component, hence inferring that removing the sodium chloride and the ammonium sulphate would make a biocompatible PP buffer.

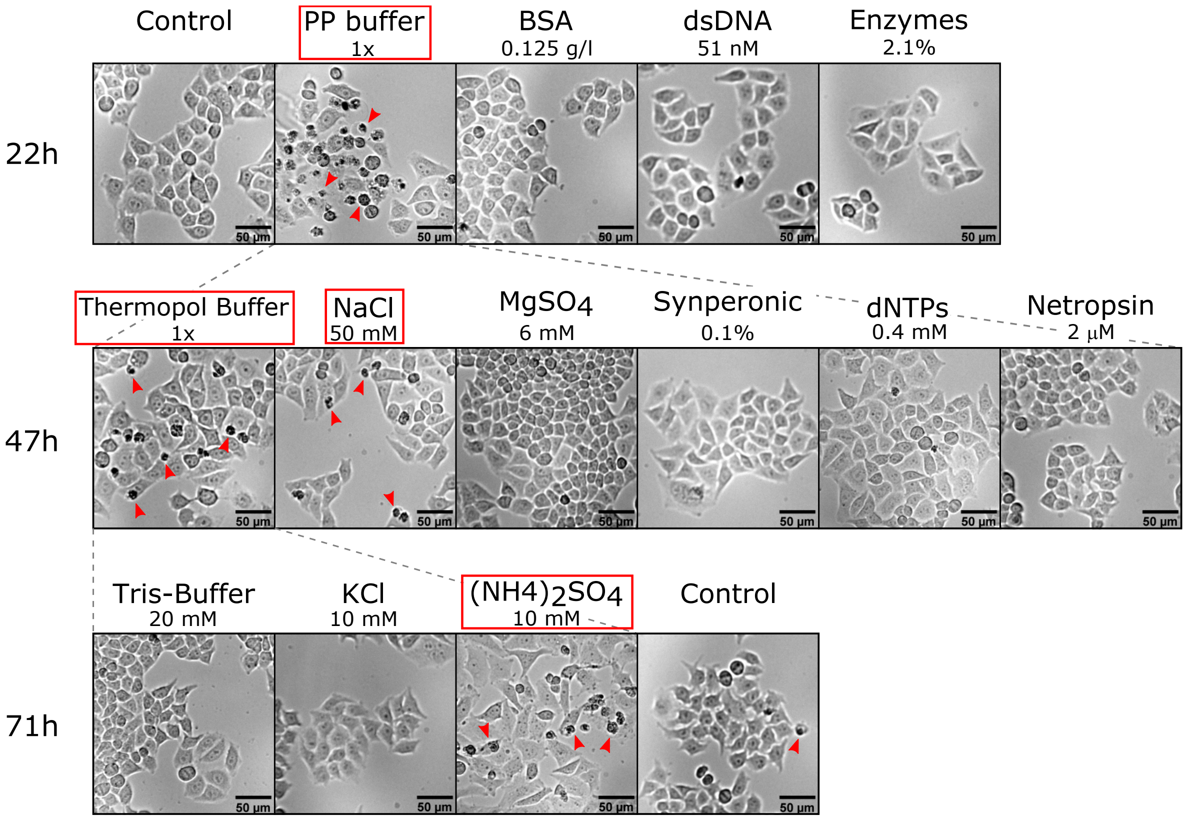


Figure S1: Screening the cytotoxic components of the PEN DNA toolbox. Bright-field images of HeLa cells after their incubation in the presence of the different components used for PEN reactions, at their respective working concentrations. The top panel shows the major components of the PEN DNA toolbox after 22 h of incubation, the middle panel the components forming the PP buffer after 47 h of incubation, and the lower panel shows the components of the Thermopol buffer after 71 h of incubation. Control shows cells incubated only in the presence of 1% Penicillin-Streptomycin and DMEM. BSA stands for Bovine Serum Albumin (New England biolabs: B9000S), and dsDNA indicates 50nM of strand $\mathbf{T}_1$ and 1nM of strand $\mathbf{A}_1$. Red arrowheads are placed to help the visualization of dead cells, which appear as black debris.

While the PP buffer resulted in being cytotoxic for cells, dithiothreitol (DTT) affected cellular integrity by causing the partial detachment of the cells from the surface (Figure S2). In particular, DTT presented a faster dynamics, observing detachment after only 4h of incubation, but also presenting a recovery stage, where cells re-attached onto the surface.

Concentrations used for standard PEN reactions (3 mM) resulted in very high levels of cellular detachment, and recovery periods longer than 2 days. Upon the reduction of DTT by one order of magnitude, recovery periods were shorter than 1 day, and the cells affected by detachment were low. Since cellular detachment affects cellular growth,[6] we included DTT as a cytotoxic components to remove (or reduce in concentration) from the DNA buffer.

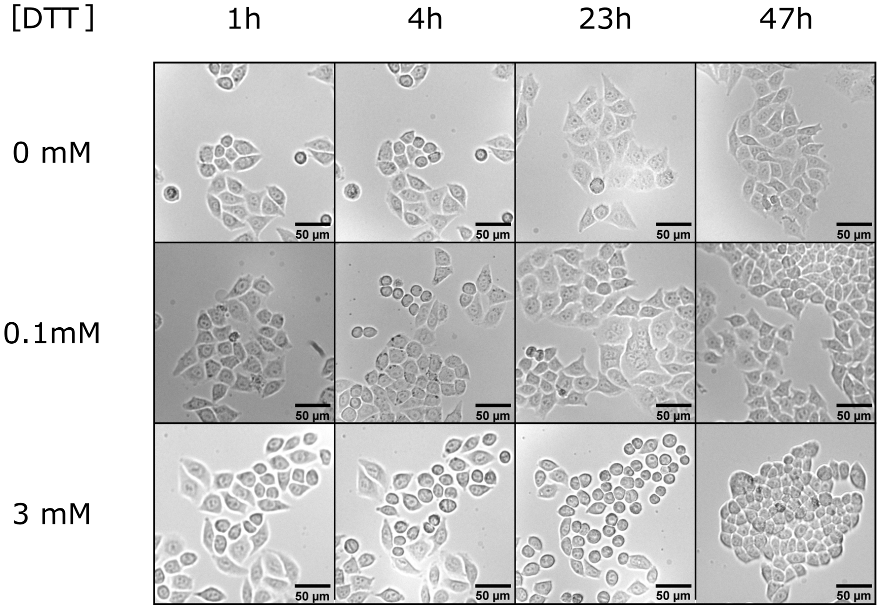


Figure S2: Dithiothreitol (DTT) causes transient cell detachment. Bright-field images at different time points of HeLa cells incubated in 0mM, 0.1 mM (used in this study) and 3 mM (standard PEN reactions) of DTT. Note: images are not always obtained in the same site.

Having identified the cytotoxic components of the PEN DNA toolbox for at least 3 days, we then tested the efficiency of the PEN reactions in the absence of these compounds. To do so, we compared the core element of the PEN reactions, the exponential amplification, at 37 °C, which is the standard temperature for growing human cells. A range in DTT concentration revealed no significant differences in the autocatalytic dynamics (Figure S3a), indicating the possibility of reducing or removing the DTT from the DNA buffer, and hence decrease cellular detachment. For the other cytotoxic component, the PP buffer, we re-designed it but without sodium chloride and ammonium sulphate, from here onwards referred as Bio-PP buffer. The autocatalytic network of species **A** yielded the expected sigmoidal curve of PEN autocatalysis reactions in both PP buffer and Bio-PP buffer (Figure S3b). An increase in the amplification onset time, $\tau$, from 40 up to 94 min, revealed a slower dynamics between the two buffers. A range of initial concentration of **A** in the autocatalytic network further coroborated this, since a 37% reduction in the rate of the autocatalytic reaction was observed between the PP buffer and the new Bio-PP buffer (Figure S3c). We attributed this reduction mainly to the absence of sodium chloride, which is essential for the efficiency of the enzymes (as specified by supplier). When DMEM was added to the Bio-PP buffer, the rate of the autocatalytic reaction reduced down to 20% (MT Figure 2a), attributing this to the sodium chloride present in the DMEM. (Note: $\tau$ and rates cannot be directly compared due to the difference in template **T** concentration used). The lower fluorescence shift of Bio-PP buffer cannot be easily explained, since it could be a lower trigger production or a difference in fluorescence behaviour of the dye (since salts concentration are important).

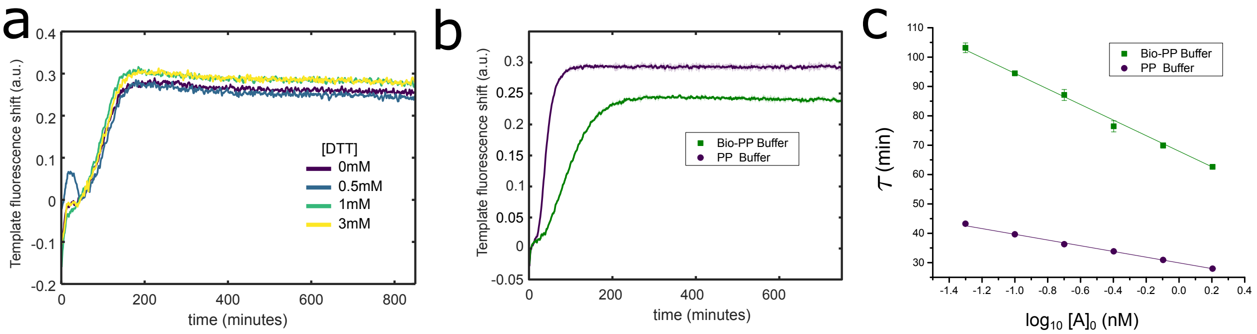


Figure S3: Dynamics of PEN reactions in the absence of cytotoxic compounds. a) Exponential amplification of strand $\mathbf{A}_1$ in PP buffer with different DTT concentrations. b) Exponential amplification of strand **A** in PP buffer or Bio-PP buffer. The shades, corresponding to the standard deviation of a triplicate experiment, are in the order of 0.01 (a.u), being in the range of the line thickness. c) Amplification onset time, $\tau$, against the decimal logarithm of the initial concentration of **A** for the autocatalytic network in PP or Bio-PP buffer. The errors correspond to the standard deviation of a triplicate experiment. Conditions: 50 nM of template, with 1 nM or 0.1 nM trigger for panel a and panel b, respectively.

Having a functional non-cytotoxic PEN reaction, the second step was to make it fully biocompatible to allow the co-existence of living cells. To do this, we had had to add the essential cell culture medium, being DMEM and 1% antibiotics. Figure S4a shows the effect on the PEN dynamics as a function of the DMEM concentration. It can be clearly observed that the higher the concentration of DMEM, the larger the delay of the amplification onset time, and the disappearance of the sigmoidal behaviour. At the highest DMEM concentration tested, 0.66x, a reduction of 5-fold in the amplification onset time was observed. Implicitly, the higher the DMEM concentration, the closer to cell culture conditions, and hence the higher biocompatibility, but the PEN dynamics were becoming too slow. For this reason, we compromised for a 0.5x DMEM concentration where PEN dynamics was only reduced by 3-fold. But, before compromising, we verified that the reduction in cell culture concentration was not adverse to cell culture viability. Figure S4b shows that cells presented normal morphology when grown in 0.5x DMEM supplemented with 0.5x PBS (to keep normal osmolarity) for at least 3 days.

To avoid unwanted contamination of the solutions containing cell culture medium, we relied on the use of 1% Penicillin-Streptomycin antibiotics. Exponential amplification of PEN reactions revealed no significant modification in either standard PP buffer or with further 0.5x DMEM in the presence or absence of 1% antibiotics (Figure S4c). To our surprise, when we tested the PP buffer with 0.5x DMEM in the absence of DTT (Figure S4d red dashed line), no autocatalytic behaviour was detected, contrary to what had been previously observed in the absence of DMEM (Figure S3a). This meant that DTT, although been cytotoxic for the cells, needed to be present. When testing a range of DTT in the final optimal biocompatible DNA buffer (Bio-PP buffer, 1% antibiotics and 0.5x DMEM), we obtained autocatalytic behaviour above 0.2 mM of DTT, with no significant changes in dynamics above this concentration. For this reason, and knowing the cytotoxicity of DTT, the Cell-DNA buffer was defined with 0.1 mM of DTT (Table S1). Biocompatibility and functionality of the Cell-DNA buffer are found in the manuscript.

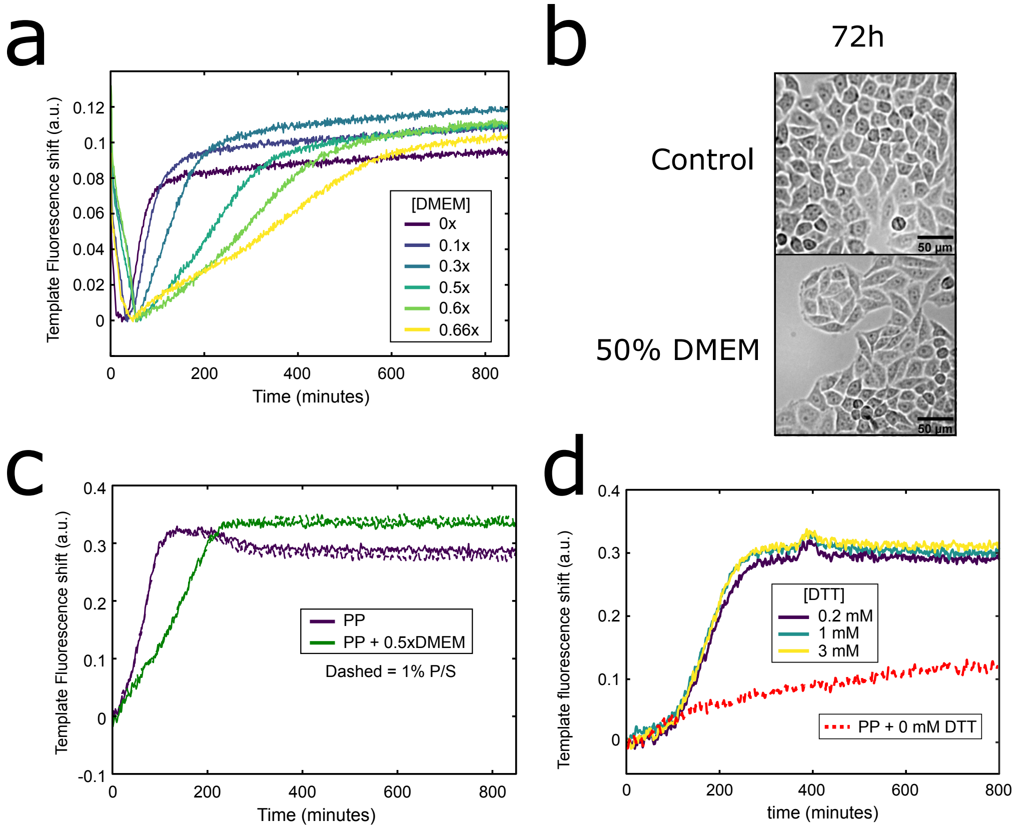


Figure S4: The PEN DNA toolbox works in a biocompatible buffer. a) Exponential amplification of strand $\mathbf{A}_2$ in standard DNA buffer (1x) supplemented with different DMEM concentrations. In this case, fluorescent shifts were calculated by normalizing by a late time point (t=995), subtracted from 1, and shifted vertically to the origin, since raw fluorescence intensities were obtained with the CFX. b) Bright-field images of Hela cells after 71h growing in 0.5x DMEM and 1% antibiotics (supplemented with 0.5x PBS). c) Exponential amplification of strand $\mathbf{A}_1$ in PP buffer (purple line) or in PP buffer with 0.5x DMEM (green line), with (dashed line) or without (solid line) 1% Penicillin-Streptomycin antibiotics. d) Exponential amplification of strand $\mathbf{A}_1$ with 0.5x DMEM in Bio-PP buffer with a range in DTT concentration (solid lines), or in PP buffer without DTT (dashed red line). Conditions: 50 nM of template, with 1 nM or 0.5 nM trigger for panel a and c, and panel d (except dashed red line which was 1 nM), respectively.

### 3 Tables S1 to S2

Table S1: Composition of the different buffers used in this work.

| **Name** | **Component** | Concentration |
|---|---|---|
| Cell-DNA buffer | | |
| | DTT | 0.1 mM |
| | BSA | 0.125 g/l |
| | DMEM | 0.5x |
| | Antibiotics | 1% |
| | Bio-PP buffer | 1X |
| Bio-PP buffer 1X | | |
| | $MgSO_4$ | 6 mM |
| | Syperonic F108 | 0.1% |
| | dNTPs | 0.4 mM |
| | Netropsin | 2 uM |
| | Bio-Thermopol Buffer | 1X |
| Bio-Thermopol buffer 1X | | |
| | Tris-HCl | 20 mM |
| | KCl | 10 mM |
| | MgSO4 | 2 mM |
| Reference Cell buffer | | |
| | Antibiotics | 1% |
| | DMEM | 99% |
| DNA buffer | | |
| | DTT | 3 mM |
| | BSA | 0.125 g/l |
| | PP buffer | 1X |
| PP buffer 1X | | |
| | $MgSO_4$ | 6 mM |
| | NaCl | 50mM |
| | Synperonic F108 | 0.1% |
| | dNTPs | 0.4 mM |
| | Netropsin | 2 uM |
| | Thermopol Buffer | 1X |
| Thermopol buffer 1X | | |
| | Tris-HCl | 20 mM |
| | KCl | 10 mM |
| | $MgSO_4$ | 2 mM |
| | $(NH_4)_2SO_4$ | 10 mM |

Table S2: DNA sequences used in this work. Phosphorothioate bonds are marked with an asterisk *, terminal phosphates with **p**, dual.bt indicates a dual biotin modification, and NH2 indicates Amino C6 modification. JOE, TR (Texas Red), AMCA, DY350, ROX, ATTO590, FAM, Dabcyl (quencher of FAM) and Cy (Cyanine) are the used fluorophores. The name of the species used in the Main Text appears in the left. The corresponding name used internally in the lab, is recalled on the right. Subscript define completely different sequences for the same node (such as templates), while superscript are slight modification of the same sequence (either 5 or 3' chemical modifications, or the addition or subtraction of a nucleotide).

| **Name** | **Sequences 5' → 3'** | **Lab name** |
|---|---|---|
| **A** | CATTCTGCGAG | Ba-A8 |
| $\mathbf{A}^1$ | **p**CATTCTGCGAG | pBa-A8 |
| **T** | JOE-*C*T*C*GCAGAATGCTCGCAGAA**p** | JOE_CBa-A8(-2)PS3 |
| $\mathbf{T}^1$ | TR-*C*T*C*GCAGAATGCTCGCAGAA**p** | TR-CBa-A8-2PS4 |
| **R** | T*T*T*TCTCGCAGAATG**p** | pTBa-A8_T4 |
| $\mathbf{R}^1$ | Cy5-T*T*T*TCTCGCAGAATG**p** | pTBa-A8_T4_Cy5 |
| **S** | Cy3.5-*C*T*T*CACCATAACCT*A*C*C*-Cy3.5 | Cy3.5_SD1*_Cy3.5 |
| random | CATCTTCATCCCATCTTCATCC | Lp*Lp* |
| **R*** | CATTCTGCGAGAAAA | pTBa-A8_T4* |
| **D** | dual.bt-*A*A*G*GTAGGTTATGGTGAAGTCTCGCAGA**p** | 2Bt_Ba-A8(-3)_2_SD1 |
| A-A | CATTCTGCGAGCATTCTGCGAG | Ba-A8_Ba-A8 |
| $\mathbf{D}^1$ | NH2-A*A*G*G*TAGGTTATGGTGAAGTCTCGCAGA**p** | NH2_Ba-A8(-3)_2_SD1 |
| $\mathbf{D}^2$ | NH2-A*A*G*G*TAGGTTATGGTGAAGTCTCGCAGAA**p** | NH2_Ba-A8(-2)_2_SD1 |
| $\mathbf{A}_1$ | CATTCTGACGAG | Ba12 |
| $\mathbf{T}_1$ | Cy3.5-*C*T*C*GTCAGAATGCTCGTCAGAA**p** | cy35-CBa12-2PS4 |
| $\mathbf{A}_2$ | CATTCAGGATCG | Be12 |
| $\mathbf{T}_2$ | Cy3.5-C*G*A*T*CCTGAATGCGATCCTGA**p** | cy35-CBe12-3PS4 |
| $F_1$ | AMCA-*C*C*AAGACUCAGCCAAGACTCAGTTTTT | AMCA-e2eT5-B |
| $F_2$ | DY350-*A*A*AAAAAAAAAAAAAA**p** | DY350-16A |
| $F_3$ | TR-*C*T*C*GCAGAATGCTCGCAGAA**p** | TR-CBa-A8-2PS4 |
| $F_4$ | ROX-CATAACACAATCACA | ROX-Th5,6:6-t |
| $F_5$ | T*T*ACTCAGCTTAGACAGATGACTCTC-ATTO590 | b2ia-AT590 |
| $F_6$ | Cy3-G*T*G*GGAGAATGAAGT-NH2 | Cy3-Li-NH2 |
| $F_7$ | Cy5-*A*A*AAAAAAAAAAAAAA**p** | Cy5-16A |
| $F_8$ | Cy5.5-A*A*A*AAACAGACUCGA**p** | Cy5.5pTdA4 |
| $ref_1$ | FAM-TCAAGTGAGTCGCAGTCCTCCTTAGCATTCGATCATAAGC-Dabcyl | Nsub |
| $ref_2$ | AGTCTGTTTCGAGTAA | inh16 |

### 4 Figures S5 to S14

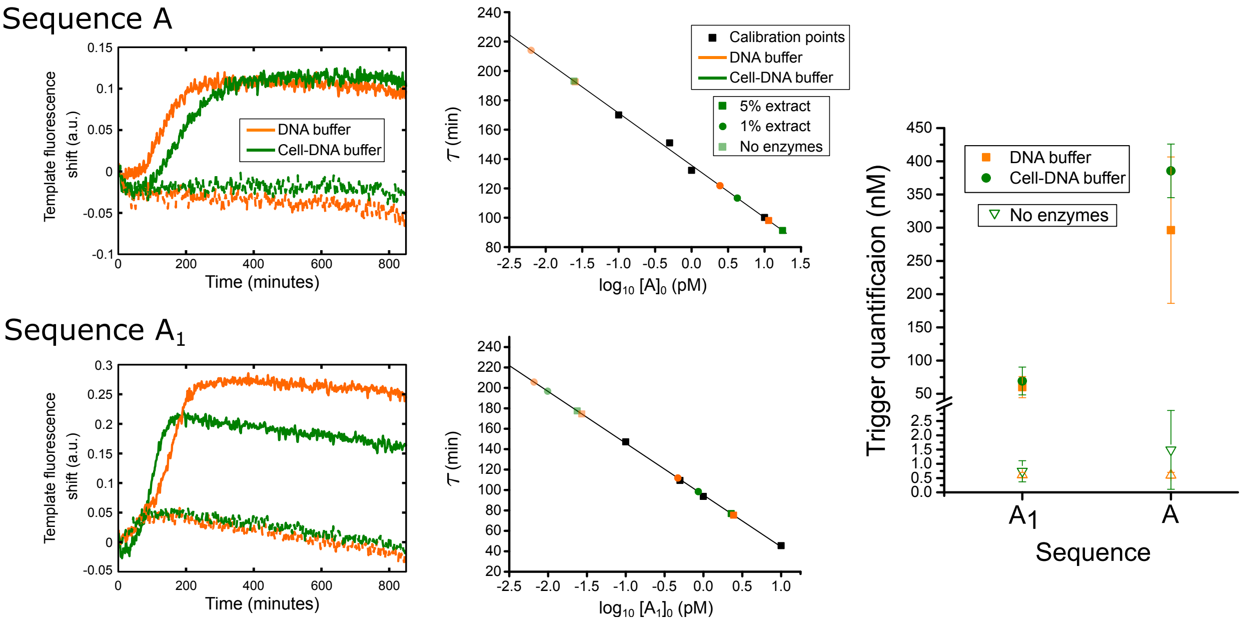


Figure S5: Isothermal quantification of trigger production at steady state for templates $\mathbf{T}^1$ and $\mathbf{T}_1$ at 37 °C. Left panel shows the exponential amplification from where solution was extracted (5% or 1%, square and circle symbol respectively) and placed into a fresh isothermal amplification, where the amplification times, $\tau$, were plotted within a trigger titration calibration curve (middle panel). Right panel shows the extracted trigger concentrations. Trigger concentration of **A** is ~10-fold greater than that of $\mathbf{A}_1$ at 37 °C, while little difference between the different buffers is observed. In the case of the negative controls (absence of enzymes), no amplification (or degradation) is expected, hence trigger concentration should be the $[\mathrm{A}]_0$ used in the left panel, 0.5 nM. Predicted trigger concentrations for the negative controls were in close agreement with the expected, showing the robustness of this technique.

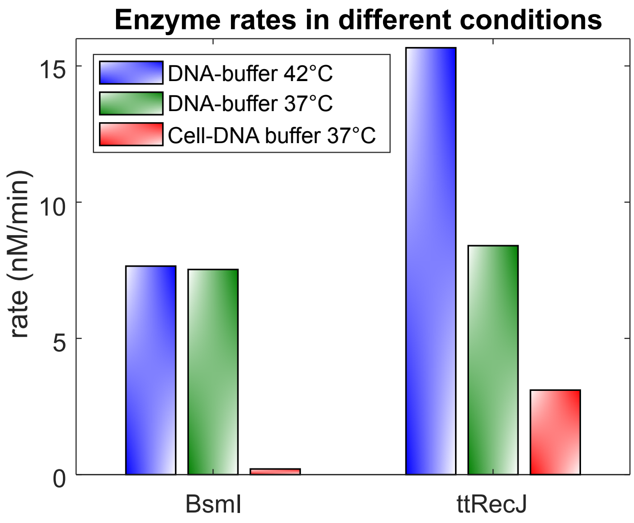


Figure S6: PEN DNA toolbox enzymatic rates measured in different buffers and temperatures. BsmI is the nicking enzymes used for the PEN reaction in the Main Text. ttRecJ is the exonuclease used in the Main text. The rates were measured at different enzyme concentrations: [BsmI] = 0.4% and [ttRecJ] = 1%.

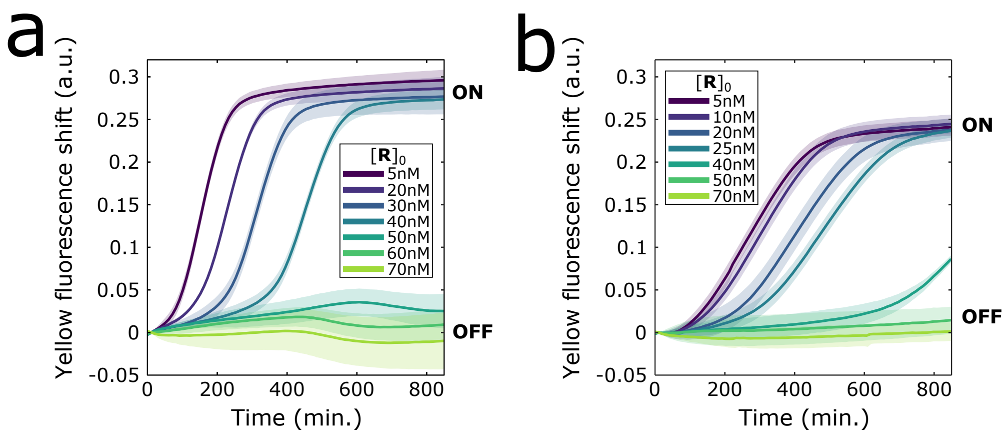


Figure S7: Bistable switch in different buffers associated to MT Figure 2d. Fluorescent shift from fluorescently-labelled **T** versus time for the bistable switch for increasing concentrations of **R** in standard DNA Buffer (a) and in Cell-DNA buffer in the absence of cells (b). Conditions are identical to MT Figure 2d.

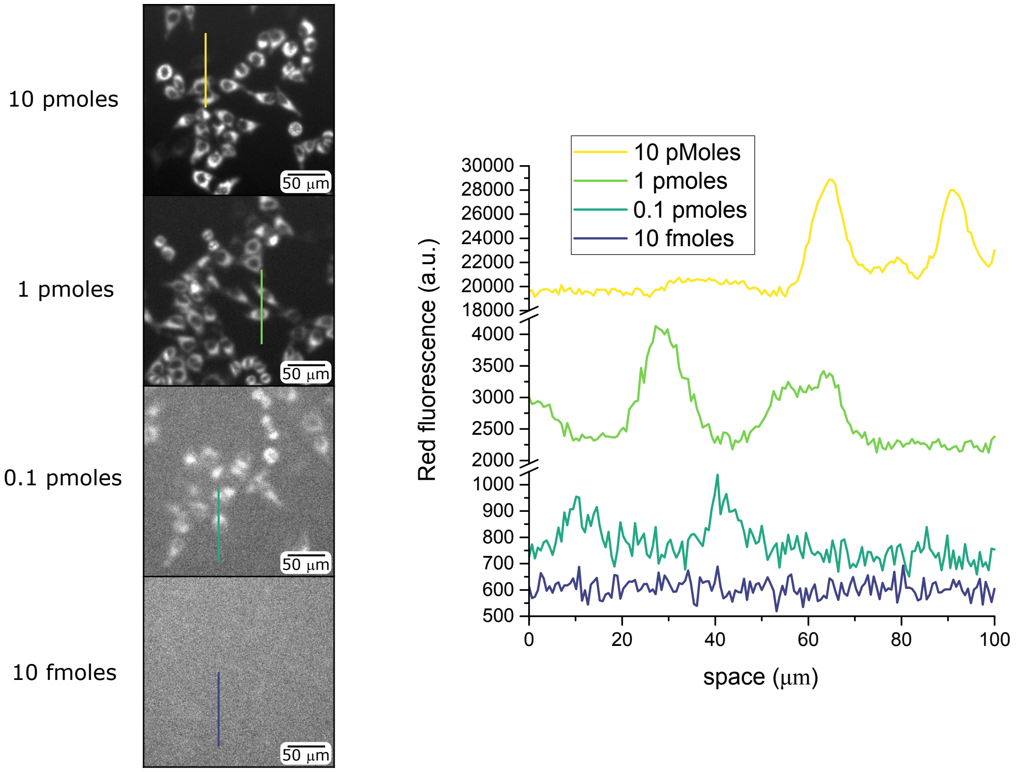


Figure S8: Internalization of Fluorescent DNA. Images of HeLa cells after 10h of incubation with different strand **S** concentrations with their respective profiles. Data shows that under 0.1 pmoles of DNA release, no DNA internalization can be distinguished from the background (no rinsing has been performed). Conditions: 2000 cells in 96 cell well plates with 100 $\mu$L of solution (1% Penicillin-Streptomycin, Strand **S**, 2.5% FBS and DMEM). Image acquisition size 648.1 $\mu m^2$.

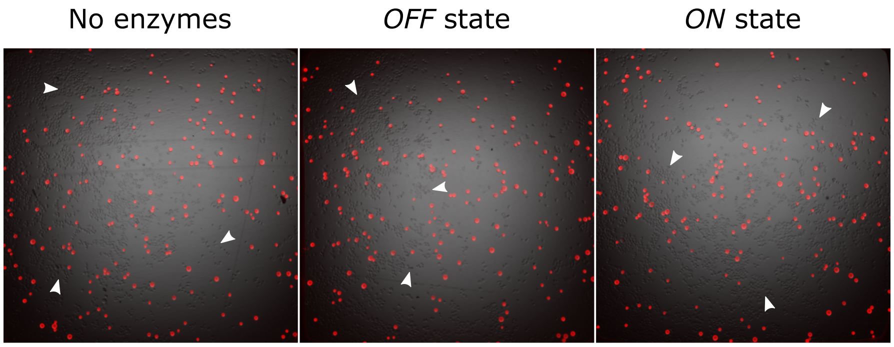


Figure S9: Homogeneity in the distribution of the cells and the conversion beads (in red). Overlapped of bright-field and red fluorescence microscopy images at the start of the experiments shown in MT Figure 3. To calculate % surface coverage of beads, 'analyze particle' routine of ImageJ was used to extract the number of particles per well, to further calculate using the average particle size provided by the manufacturer, 34 $\mu$m. With 161 particles per well ($n = 18$), a 2.6% (with a standard deviation of 0.4%) surface coverage by the beads was predicted, with a total releasing capacity of $\sim$0.8 pmoles of Strand **S**, concentration which should be visible according to Figure S8. Arrowheads are placed to help the visualization of cells.

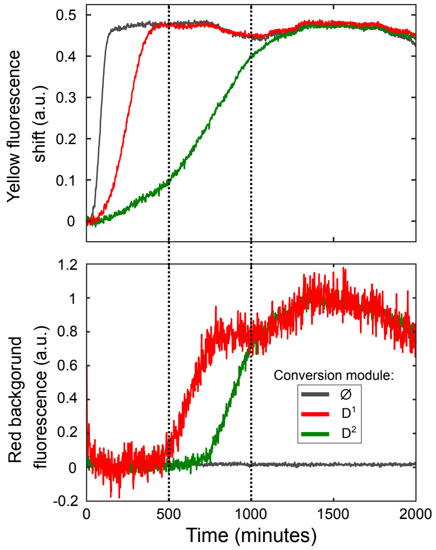


Figure S10: Controlling the release dynamics of strand **S**. As previously showed by Montagne *et al.*,[7] strand **A** has a higher affinity to **R** than to **T** because the region of complementarity between **R** and **A** is larger than that between **T** and **A** (Table S2). This difference makes the hybridization of **A**:**R** more thermodynamically stable, and hence predominate, over the **A**:**T** dsDNA. In the same manner, the conversion module can be tweaked to favour the hybridization **A**:**T** over **A**:**D**. When the conversion module has the same region of complementarity as that of **T** ($\mathbf{D}^2$, green line), the conversion module slows the exponential amplification by 9-fold. We account this to the fact that $\mathbf{D}^1$ and **T** both share **A** at low concentrations, which is further corroborated by the fact that both modules finish at relative similar times (∼1000 min). On the other hand, when using a conversion module which has a lower region of complementarity than that of **T** ($\mathbf{D}^1$, red line), the autocatalytic module is only slowed by 3-fold. In this case, we don't attributed this to the activity of the conversion module, since no release of the load is observed until steady state is reached (∼500 min). Hence, by reducing the affinity of **A** onto **D** than that to **T**, the autocatalytic and the conversion modules finish before, and assures the conversion module is only active when steady state has been reached. $\phi$ is in the absence of conversion module. The black dashed lines are guides to the eye. Conditions: DNA buffer with 50 nM of **T**, 0.5 nM of $[\mathrm{A}]_0$, and 20 nM of conversion module $\mathbf{D}^1$ or $\mathbf{D}^2$ loaded with 15 nM of strand **S**. Note: we expect no great difference in dynamics due to the presence of dual biotin (**D**) or Amino C6 ($\mathbf{D}^1$ and $\mathbf{D}^2$).

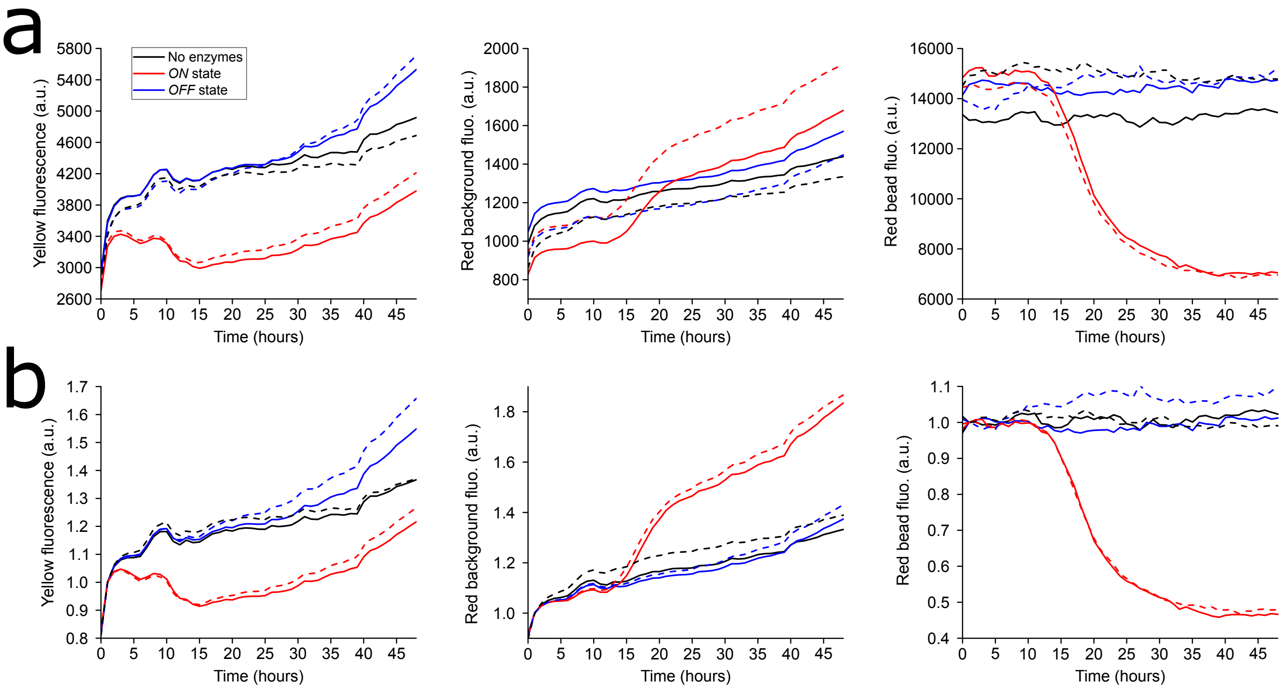


Figure S11: Raw (a) and background-corrected data (b) for one subset of the data presented in MT Figure 3. a) Fluorescence versus time plots displaying the raw data of the internalization switch in the *ON* (red line) and *OFF* state (blue line), and in the absence of enzymes (black line) for 2 replicates (full and dashed line) performed the same day. b) Same fluorescence versus time plots as in panel where each curve has been divided by the florescence value at an early time point. We attribute the higher standard error observed in MT Figure 3 compared to panel b, up to 2 orders of magnitude, due to human derived experimental differences between the different experimental days. Conditions are identical to MT Figure 3.

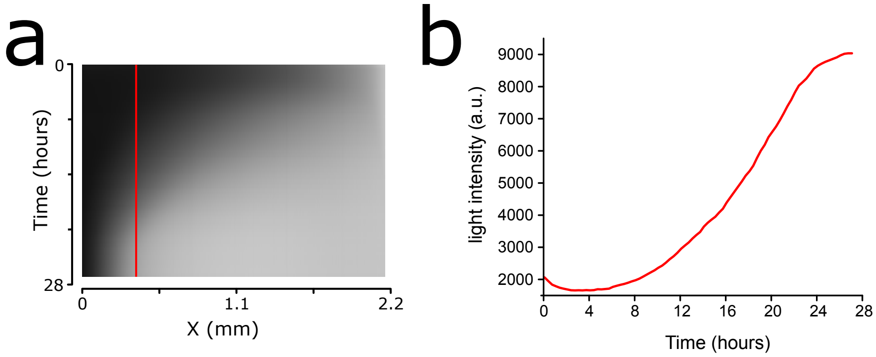


Figure S12: Evaporation dynamics from 384 cell well plates during experimental procedures at 37 °C. a) Kymograph tracking the evaporation of 5 $\mu$L of DMEM medium in a 384 cell well plate over time. Note that 5 $\mu$L is not sufficient to wet the entire bottom of the well (spatial heterogeneity). b) Profile depicted in panel a. As the DMEM medium evaporates, the light intensity of the well increases until it reaches saturation, where no more changes in the light path occur. An evaporation rate of 4.44 $\mu$L/day is foreseen to occur in our setup, giving rise to a maximal total loss of 27% during 2 days. As volume in the 384 cell well plate decreases. the concentration of the fluorescently-labelled **T** increases, and so does its fluorescence signal. Since the yellow fluorescence shift graphs depict the quenching phenomenon of the fluorophore, the increase in the fluorescence due to evaporation transduced into a decrease in the yellow fluorescence shift (observed in MT Figure 3 and 4).

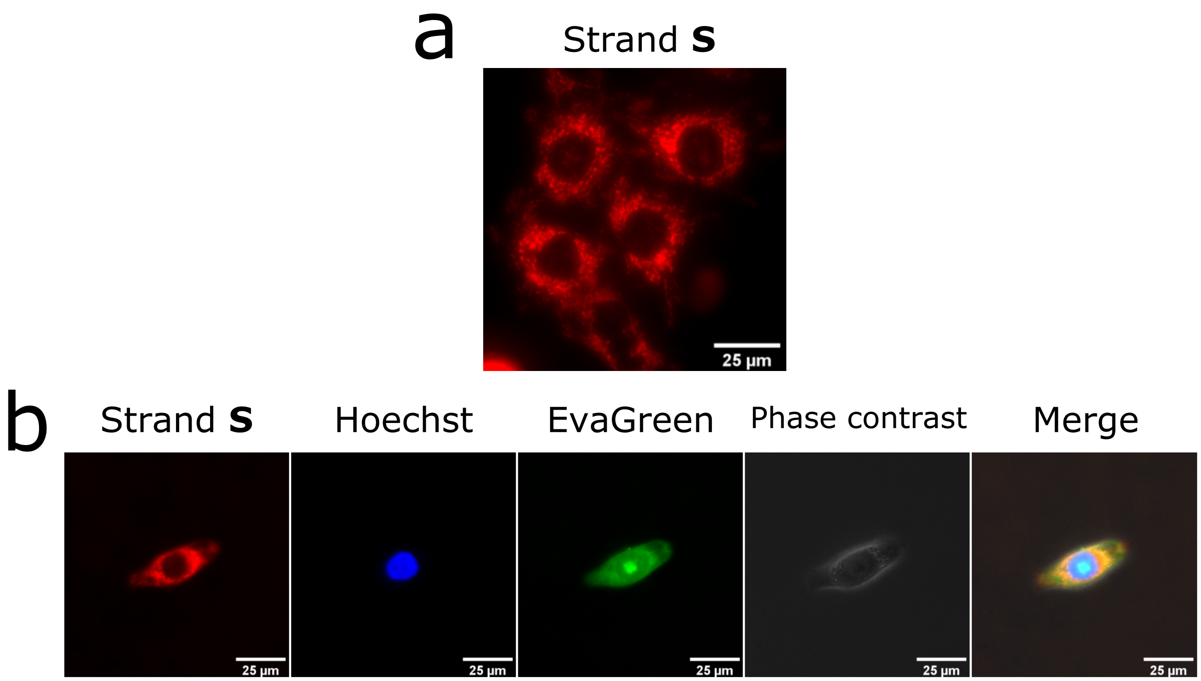


Figure S13: Strand **S** internalizes into dot and rod-like compartments. a) Dried cell images showing the spatial distribution of strand **S** into rod-like compartments. b) Imaging of living cells in the presence of the strand **S**, a nuclear staining (Hoechst), EvaGreen (which resulted in the staining of the membrane and partially of the inner nucleus) and phase contrast. The merge reveals strand **S** is not evenly distributed within the cytoplasm, but rather focused closely to the nucleus of the cell. Images obtained with a 40X objective. Image acquisition size 166.4 $\mu m^2$ and 332.8 $\mu m^2$ for the top and bottom panel, respectively.

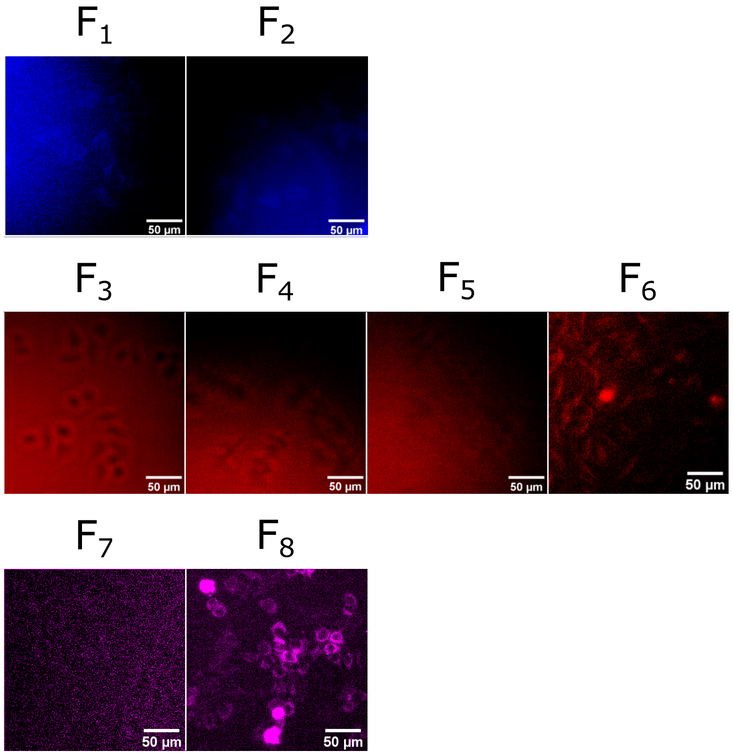


Figure S14: Detection capabilities of other standard DNA-fluorescent markers. Due to different molecular structures, some standard DNA-fluorescent markers may not be internalized or do not show fluorescence upon internalization. We screened the detection capabilities of different fluorescently labelled strands ($F_1$-$F_8$) after 24h of incubation in the presence of living HeLa cells. In addition to the marker used in this manuscript (Cyanine3.5), we only observed internalized fluorescence for Cyanine5.5 markers ($F_8$). We observed difference in behaviours within the same family of fluorescent markers, since we didn't detect internalization for Cyanine 3 ($F_6$) or Cyanine 5 ($F_7$), which present very small structural differences. Conditions: 1 pMoles of each strand was added to a final volume of 50 ul or 100ul for 384 or 96 cell well plates with 3200 or 1600 cells per well, respectively. Blue, red and far-red filters were used for the top, middle and bottom panel, respectively. Note: The internalization of the different fluorophores was tested with different sequences bearing or not PTO, and terminal-phosphate modification, which could influence also the internalization yield.

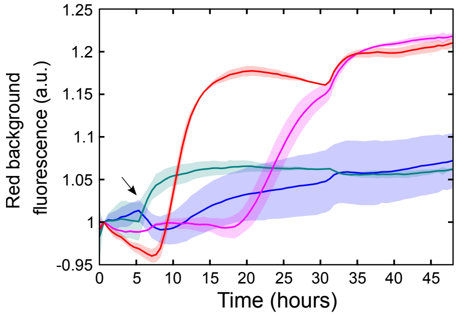


Figure S15: Release of the load **S** into the medium by the response of the internalization switch in MT Figure 4a and b. Fluorescence versus time plots displaying the dynamics release into the solution in the *ON* (red line) and *OFF* (blue line) states, as well as *OFF* states where a ssDNA, either $\mathbf{R}^*$ (pink) or a random sequence (teal), which were introduced at t = 6 h (indicated with an arrow). The bump at $t = 6$ h is an artifact resulting from the addition of the ssDNA. Conditions are identical to MT Figure 4a and b.

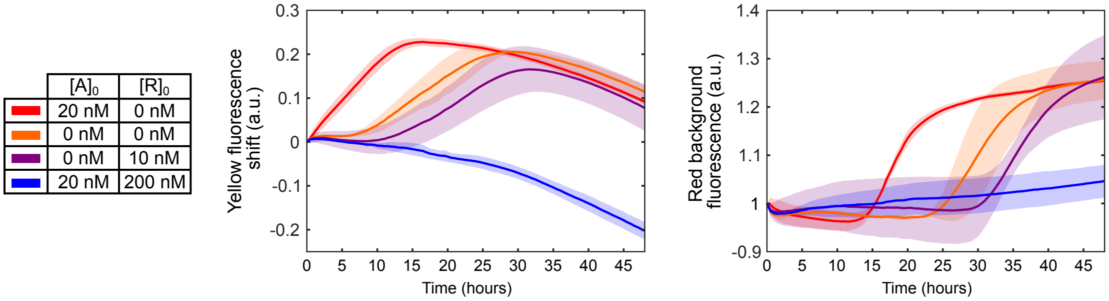


Figure S16: Pre-programmed temporal release of the load **S** into the medium. Release of the load **S** into the medium by the response of the internalization switch in MT Figure 4c and d. Fluorescence versus time plots displaying the dynamics of production of **A** (left) and the release of **S** into the solution (right) in the presence of cells with different initial conditions. The *ON* and the *OFF* states are represented by the red and the blue conditions, respectively, while in the absence of initial trigger and in the presence of low **R** are represented by the orange and the purple conditions, respectively. A delay on the exponential amplification is observed due to the absence of initial trigger, compared to the *ON* state. This delay is further increased in the presence of 10 nM of **R**, where at high concentration of **R** (*OFF* state), the exponential amplification never occurs. The delay on the exponential amplification is transferred into a systematic delay of the release of **S** into the solution. Conditions are identical to MT Figure 4c and d.

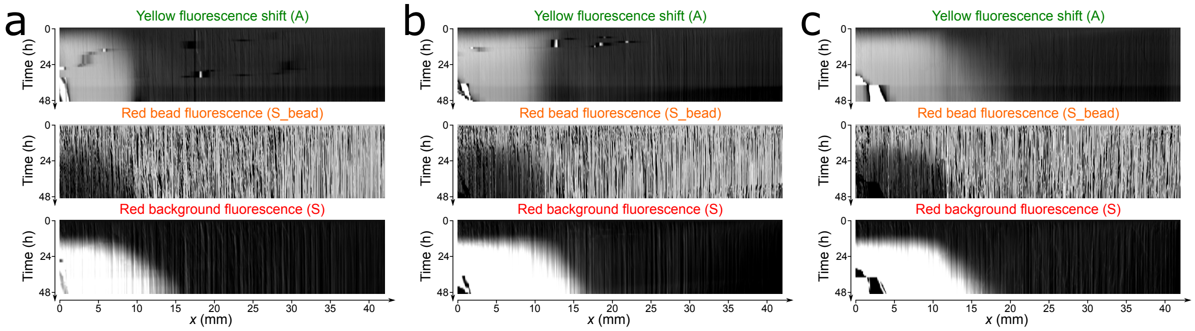


Figure S17: Spatiotemporal dynamics of the internalization switch in the absence of cells. a) Kymographs representing the behaviour of the autocatalyst, yellow fluorescence shift (**A**), the Red bead fluorescence (S_bead), and of the Red background fluorescence (**S**) with a gradient of repressor (**R**) performed with 65 $\mu$L as the injecting volume. b) idem to panel a but using 55 $\mu$L, 10 $\mu$M of **R** and pipetting back and forth 4 times instead of 5 to generate the gradient. c) idem to panel b but using 2 $\mu$M of **R**.

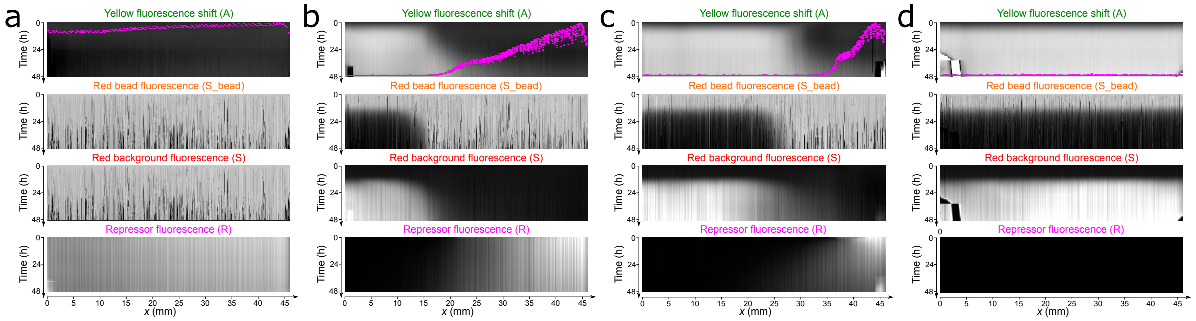

Figure S18: Spatiotemporal dynamics of the different repressor (**R**) patterns shown if MT Figure 5 f-g. a) Kymographs representing the behaviour of the autocatalyst, yellow fluorescence shift (**A**), the Red bead fluorescence (S_bead), the Red background fluorescence (**S**) and the Repressor fluorescence (**R**) starting with a high concentration of **R** (*OFF* state). b) Kymographs showing the spatiotemporal behaviour in the presence of a gradient of **R** performed using 55 $\mu$l as the injecting volume. c) Idem to panel b but in the presence of a gradient of **R** performed using 25 $\mu$l as the injecting volume. d) Idem to panel a but in the absence of **R** (*ON* state). The magenta line over-imposing the yellow fluorescence shift kymographs represents the initial **R** pattern (t= 0 h). Panels a and d show the raw fluorescence of **R**, while in panels b and c the fluorescence has been normalized between 0 and 1. Conditions are identical to MT Figure 5.

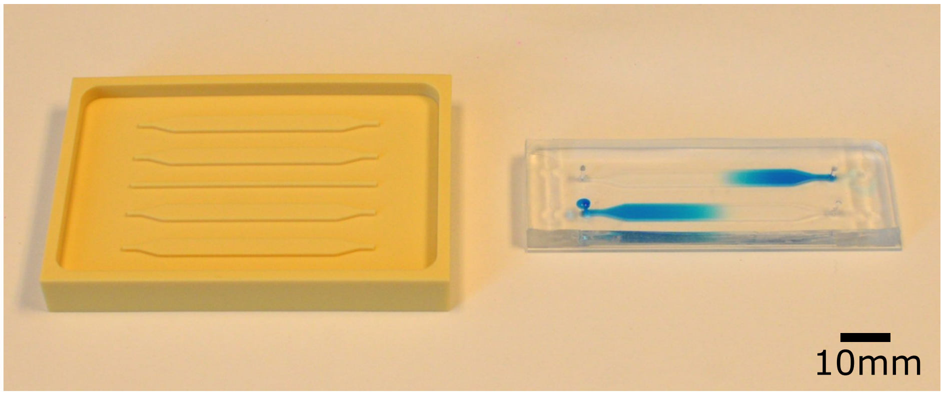


Figure S19: Mold and millifluidic device used for experiments in MT Fig. 5. Left: reusable polyvinyl chloride (PVC) mold fabricated using a milling machine. Right: millifluidic device made of molding poly(dimethylsiloxane) and sticking it on a glass slide. The channels are 50 mm long, 3 mm wide and 1 mm high. Here, a 2-channel device with two gradients of methylene blue dye is shown.

### 5 Video S1

Video S1: Timelapse showing the internalization of fluorescently-labelled DNA (strand **S**) by HeLa cells in under 4h. Conditions: 1 pmoles experiment of Figure S8.

### References


1. Zadorin, A. S., Rondelez, Y., Gines, G., Dilhas, V., Urtel, G., Zambrano, A., Galas, J. C., and Estevez-Torres, A. (2017) Synthesis and materialization of a reaction-diffusion French flag pattern. *Nature Chemistry 9*, 990–996.

2. Montagne, K., Plasson, R., Sakai, Y., Fujii, T., and Rondelez, Y. (2011) Programming an in vitro DNA oscillator using a molecular networking strategy. *Mol Syst Biol 7*, 466, 10.1038/msb.2010.120.

3. Baccouche, A., Montagne, K., Padirac, A., Fujii, T., and Rondelez, Y. (2014) Dynamic DNA-toolbox reaction circuits: A walkthrough. *Methods 67*, 234–249.

4. Urtel, G., Van Der Hofstadt, M., Galas, J. C., and Estevez-Torres, A. (2019) REXPAR: An Isothermal Amplification Scheme That Is Robust to Autocatalytic Parasites. *Biochemistry 58*, 2675–2681.

5. Owczarzy, R., Moreira, B. G., You, Y., Behlke, M. A., and Walder, J. A. (2008) Predicting Stability of DNA Duplexes in Solutions Containing Magnesium and Monovalent Cations. *Biochemistry 47*, 5336–5353.

6. Held, K. D., Sylvester, F. C., Hopcia, K. L., and Biaglow, J. E. (1996) Role of Fenton Chemistry in Thiol-Induced Toxicity and Apoptosis. *Radiation Research 145*, 542–553.

7. Montagne, K., Gines, G., Fujii, T., and Rondelez, Y. (2016) Boosting functionality of synthetic DNA circuits with tailored deactivation. *Nature Communications 7*, 13474.
